# Decoding the Microbiome-Metabolome Nexus: A Systematic Benchmark of Integrative Strategies

**DOI:** 10.1101/2024.01.26.577441

**Authors:** Loïc Mangnier, Antoine Bodein, Margaux Mariaz, Marie-Pier Scott-Boyer, Alban Mathieu, Neerja Vashist, Matthew S. Bramble, Arnaud Droit

**Affiliations:** Centre de Recherche du CHU de Québec-Université, Laval, Université Laval, G1V 4G2, Québec, Canada; Department of Pathology and Laboratory Medicine, UCLA, USA; Center for Genetic Medicine Research, Children’s Research Institute, Children’s National Hospital, Washington, DC, USA; Department of Genomics and Precision Medicine, The George Washington University of Medicine and Health Sciences, Washington, DC, USA; Département de Médecine Moléculaire, G1V 0A6, Québec, Canada

**Keywords:** multi-omics, metagenomics, metabolomics, benchmark, statistical methods

## Abstract

**Background:** The exponential growth of high-throughput sequencing technologies was an incredible opportunity for researchers to combine various -omics within computational frameworks. Among these, metagenomics and metabolomics data have gained an increasing interest due to their involvement in many complex diseases. However, currently, no standard seems to emerge for jointly integrating both microbiome and metabolome datasets within statistical models.

**Results:** Thus, in this paper we comprehensively benchmarked nineteen different integrative methods to untangle the complex relationships between microorganisms and metabolites. Methods evaluated in this paper cover most of the researcher’s goals such as global associations, data summarization, individual associations, and feature selection. Through an extensive and realistic simulation we identified best methods across questions commonly encountered by researchers. We applied the most promising methods in an application to real gut microbial datasets, unraveling complementary biological processes involved between the two omics. We also provided practical guidelines for practitioners tailored to specific scientific questions and data types.

**Conclusion:** In summary, our work paves the way toward establishing research standards when mutually analyzing metagenomics and metabolomics data, building foundations for future methodological developments.

## Background

The recent development of high-throughput sequencing technologies has permitted the generation of omics data at an exponential scale. Combining different high dimensional biological datasets within computational models represents a wonderful opportunity for researchers to better understand the underlying biological mechanisms involved in diseases^1^. Microorganism-metabolite interactions have gained an increasing interest due to their potential involvement in a large set of traits. It has been demonstrated that shifts in the microbiome-metabolome interactions have important implications on individual health ^2,3^. Indeed, recent studies for cardio-metabolic diseases ^4^ or autism spectrum disorders ^5^ have shown that the pathoetiology could be explained by a complex interplay between microbes and host metabolites ^6^ or by disruptions in the microbiota-derived metabolite processes ^7^. Thus, efficiently incorporating microbiome and metabolome data within statistical frameworks offers critical insights on the complex architecture occurring between diet or lifestyle factors on the microbe-metabolite recomposition and remains an important challenge to adequately identify biological pathways, with interesting avenues for clinical applications ^8^. However, the tremendous amount of available statistical models makes the choice of the right method a daunting task for many researchers. Typical research questions include either the inference of interactions occurring between species and metabolites or in predicting one omics borrowing the biological information contained in the other omics ^9,10^. On the one hand, the inference of interactions refers to understanding and identifying the complex relationships and dependencies between microbial communities and the metabolites they interact with. This inference aims to uncover causal relationships, dependencies, and mechanisms that drive microbial-metabolite interactions in biological systems. On the other hand, other multi-omics integration techniques use multiple types of omics data together to make predictions about biological phenomena. Advanced machine learning or deep learning methods, including multi-omics data are often employed to tackle these prediction tasks effectively. However, while models have been shown to be powerful to predict metabolome levels based on microbial compositions, they lack the mechanistic interpretation needed for building new sets of biological hypotheses.

Thus, to elucidate the complex entanglement between microorganisms and metabolites, several strategies can be exploited, each exhibiting associations occurring at different scales. Indeed, consistent with a recent report from Deek et al., 2024 ^11^, traditional workflows include distinct types of analysis, addressing complementary biological questions^2^. Briefly, common pipelines include the detection of *global associations*, *data summarization*, *individual associations*, and identification *of core features*. Firstly, researchers are often interested in determining whether a global association is occurring between the two omics. For example, one can look for a global change in metabolome levels due to a microbial recomposition induced by a specific diet or lifestyle^2^. Testing for global associations can be performed using multivariate methods such as the Procrustes analysis ^12^, the Mantel test ^13^, or the multivariate microbiome regression-based kernel association test (MMiRKAT) ^14^. This step frequently precedes the application of subsequent analyses such as *data summarization* methods or the *identification of core features* ^2^. Then, following approaches used for single omics, a common research objective is to summarize information contents in the two omics, facilitating the visualization and interpretation of large-scale biological data ^1^. The presence of two types of omics allows the exploitation of the intra- and inter-correlation existing between features of the two datasets. Application of data summarization methods including Canonical Correlation Analysis (CCA) ^15^, Partial Least Square (PLS) ^16^, Redundancy Analysis (RDA) ^17^ or more recently the Multi-Omics Factor Analysis (MOFA2) ^18^ is a major step to uncover features explaining a substantial proportion of data variability. Indeed, applications of data summarization methods have allowed the identification of taxonomic groups or metabolites involved in Type 2 diabetes ^19^. However, both *global association* and *data summarization* methods fail to provide individual relationships between one or several microorganisms and metabolites. This aspect remains central to highlight core features involved in a particular biological context. As an illustration, methods for detecting individual associations may prove relevant for the identification of bacterial genus associated with dietary-impacted metabolites ^2^. One strategy is to compute a measure of association between each metabolite-microbiota pair, using either a correlation or a regression model. Although easily implementable and interpretable, these approaches suffer from lack of power induced by the number of models fitted, limiting result transferability. An alternative way is to employ univariate or multivariate feature selection methods to adequately identify key actors at a large scale. The least absolute shrinkage and selection operator (LASSO) is a method initially developed to improve predictability while proceeding to feature selection ^20^. Indeed, the LASSO can set coefficients to zero, facilitating identification of core features. Consistently with this idea, sparse CCA (sCCA) ^21^ or sparse Partial Least Square (sPLS) ^22^ are multivariate penalized methods summarizing data variability while proceeding to feature selection. However, due to the complex structure of both microbiome and metabolome data, standard methods fall short of providing unbiased associations, limiting the biological interpretation of results.

By using either functional gene screening or sequencing analysis, metagenomics sequencing technologies can provide a general picture of the microbial composition in a given environment ^23^. However, metagenomics data highlight hard-to-analyze characteristics^24,25^. Indeed, it is now globally accepted that microbiome datasets are over-dispersed, zero-inflated, highly correlated, and compositional. Without adequate transformation the inherent compositionality of the data makes the application of standard methods incorrect, leading to inconsistent results ^24–26^. Metabolomics is the comprehensive study of all metabolites within a biological system, using analytical methods like LC-MS-based metabolomics to measure metabolite levels and small molecular mass compounds under specific biological conditions ^27^. This field is divided into untargeted metabolomics, which covers a broad range of metabolites, and targeted metabolomics, which concentrates on specific metabolites. Metabolomics data shares some features with metagenomics data, exhibiting over-dispersion and high correlation structures ^26^. Thus, interpreting metabolomics data using statistical analysis requires a particular attention for explicitly accounting for the data structure needed for robust biological interpretations. Moreover, integrating metabolomics data into pathway mapping, with other omics data, such as metagenomics, facilitates the discovery of biomarkers or metabolic signatures involved in diseases. Tools like MetaboAnalyst provide resources for these analyses ^28^.

Thus, combining these two omics together within statistical frameworks requires particular attention. Approaches to deal with compositional data either as an outcome or explanatory variable have already been proposed ^25,29,30^, covering applications of global association methods, data summarization, individual associations, or identification of core features. Conventional strategies include utilization of standard methods after suitable data transformations or purely compositional approaches ^31–34^. Subsequently, determining which strategy is the best depending on the research question remains an open problem with major implications for practitioners.

Despite recent studies proposing to evaluate statistical methods jointly integrating microbiome and metabolome data, efforts were only made either on literature reviews or on real-data applications, where the ground truth is missing. This aspect is pivotal for performing agnostic comparisons required for an unbiased evaluation ^11^. This systematic evaluation is still needed for highlighting best practices required for result interpretability and reproducibility. To our knowledge there is no systematic benchmark of integrative methods to link microbiome with metabolome datasets; constantly pushing researchers to make their choice without any robust comparison. For example, researchers often consider centered log-ratio or isometric log-ratio transformations when normalizing microbiome data with no clear insights on which method performs the best. Thus, in this paper, we comprehensively compared nineteen different integrative methods to decipher the complex entanglement occurring between microorganisms and metabolites. Methods and strategies were selected based on a recent report ^11^, covering most of the researcher’s aims, such as *global associations*, *data summarization*, *individual associations*, or *feature selection* (Figure 1). Our extensive simulation studies provide insightful lessons on the strengths and limits of methods commonly encountered in practice. Then, we applied best methods to real data on the gut microbiome and metabolome for Konzo disease ^35^, highlighting a complex architecture between the two omics occurring at different scales. Finally, we provide general and specific recommendations for researchers depending on the data at-hand and the research aims, while pinpointing avenues for future methodological developments (Tables 1-2). A complete user guide with all the related code is provided, facilitating method applications to other contexts, promoting scientific replicability and reproducibility.

**Figure 1:**
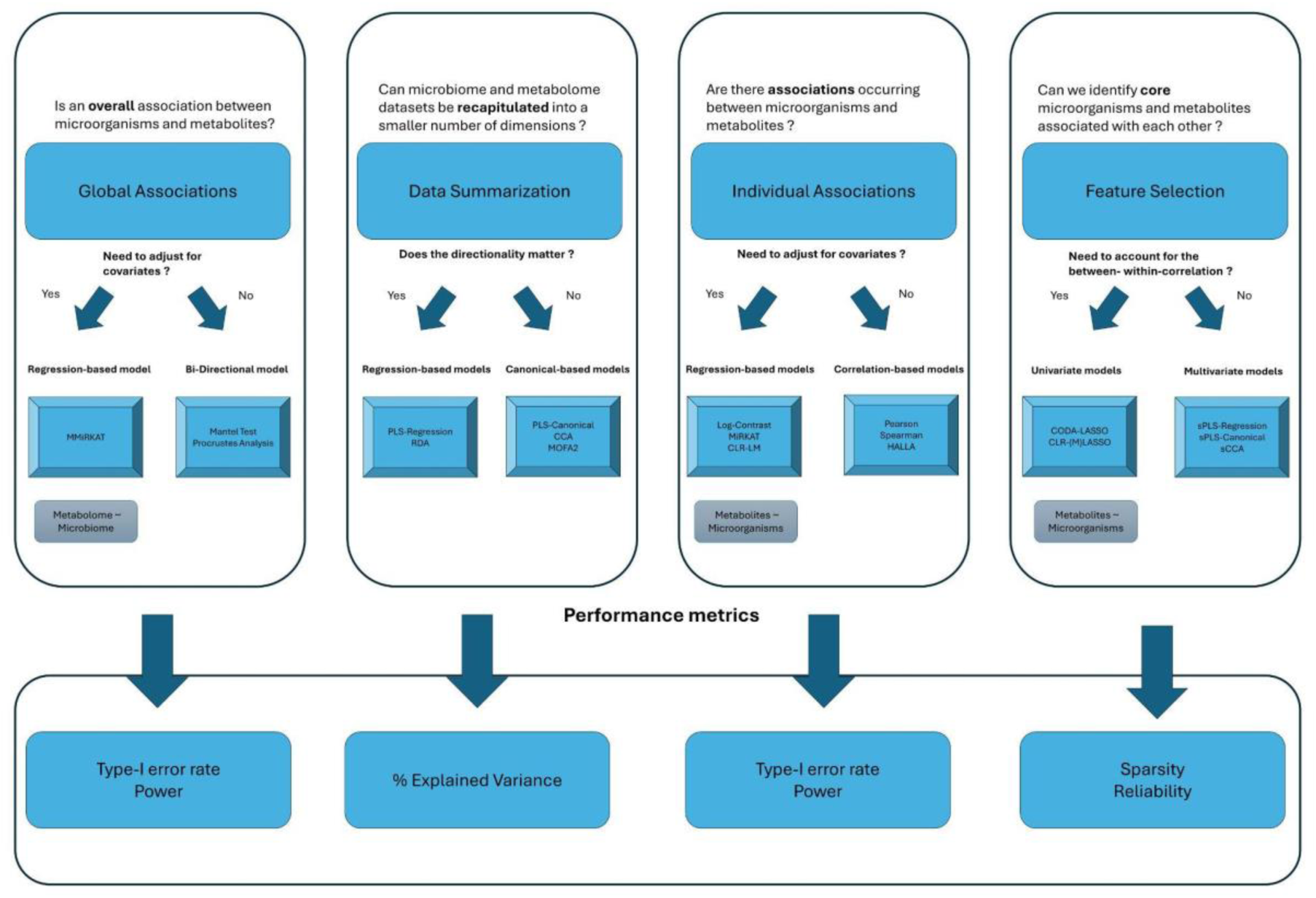
Flowchart of the integrative methods selected across the benchmark and their related performance metrics depending on the research question.

**Table 1:** Summary of best methods depending on the research question.

| Scientific Question | Research Aim | Best Method | Pros | Cons | Implementation |
| --- | --- | --- | --- | --- | --- |
| Is there any relationship between microorganisms and metabolites at a global level? | Global associations | Mantel Test | Robust to data normalization and distance kernels<br>Computationally efficient | Cannot incorporate covariables to adjust for | Vegan R package |
| Are microbiome and metabolome datasets summarizable through a limited number of components? | Data summarization | RDA | Robust to data normalization and distance kernels | Can only capture linear effects<br>Sensible to directionality | Vegan R package |
| Can we identify associations between metabolites and species? | Individual associations | MiRKAT | Allow adjustment for covariates<br>Robust to microbiome normalization | Limited to few families of generalized linear models | MiRKAT R package |
| Can we identify core microorganisms and metabolites? | Feature selection (univariate) | CODA-LASSO (compositional covariates) | Compositional and sub-compositional coherent<br>No need to data transformation<br>Allow adjustment for covariates | Limited to few families of generalized linear models<br>Long running times in high-dimensional problems | Coda4microbiome R package |
|  | Feature selection (multivariate) | sPLS | Flexible framework<br>Efficiently account for within- between- correlation<br>Robust to directionality | Tuning parameters | mixOmics R package |

**Table 2:** Overview of avenues for future methodological developments to jointly analyze metagenomics and metabolomics data. Three research objectives with their corresponding methods were identified as important avenues for future methodological developments.

| Objective | Methods and Limits | Methodological avenues |
| --- | --- | --- |
| Data normalization | - **CLR**: provides still-correlated features in the original space<br>- **ILR; alpha**: are “black-box” transformations providing uncorrelated features in a restricted space | Data normalizations providing uncorrelated features in the original space for facilitating result interpretation. |
| Mechanistic interpretation | - **Log-contrast; CODA-LASSO**: unable to provide a mechanistic view of modifications between microbiome and metabolome data | Applications of differential integrative strategies such as DIABLO <sup>51</sup> or Network-based model to jointly study the modifications of microorganism and metabolite co-occurrence networks <sup>52</sup> |
| Feature selection | - **CODA-LASSO; sPLS**: lack of sparse solutions | Extension to knockoff framework <sup>47</sup> or stability selection <sup>48</sup> for improving feature selection performances |

## Results

### SIMULATION SETUP AND BENCHMARKED METHODS

To adequately simulate realistic simulation scenarios, we generated synthetic microbiome and metabolome datasets mimicking complex data structures and relationships from three real metagenomics-metabolomics studies. Globally, datasets were chosen to exhibit different structures and complexity, covering most scenarios commonly encountered in practice. See Figure 2 and the Method section (Simulation setup) for additional details on the simulation setting. We therefore compared nineteen integrative methods for inferring the link between microorganisms and metabolites occurring at different biological scales (Figure 1). Methods were presented as follows. First, In the *global associations* subsection we compared the Procrustes analysis, Mantel test, and MMiRKAT with respect to the Type-I error rate and power. Second, In the *data summarization* subsection we evaluated four different models including CCA, PLS, RDA and MOFA2, regarding their capability to recapitulate data variability across latent factors. Third, in the *individual associations* subsection we compared four strategies for performing regression-based approaches between compositional covariates and metabolites, the clr-linear model, the log-contrast, HALLA, and MiRKAT. Approaches were evaluated based on the Type-I error rate and power. Fourth, in subsections *univariate feature-selection for compositional predictors* and *multivariate feature-selection* we compared approaches for identifying core microbes and metabolites, leveraging both univariate and multivariate feature selection strategies. For univariate frameworks, depending on the nature of the response, several models were considered. Indeed, when microorganisms are the explanatory variables, we compared three approaches, the clr-LASSO, the clr-MLASSO and CODA-LASSO ^30^. Nonetheless, for multivariate feature selection models, we considered sCCA and sPLS. Approaches were evaluated based on sparsity and reliability. Whenever suitable, we evaluated the impact of data normalizations or distance kernels on method performance. Details on the methods, data transformations, and performance metrics were provided in the Methods section. Finally, to highlight complementary biological insights provided by methods, best approaches were illustrated in the *real-data application* subsection, exploiting metagenomics and metabolomics data from Konzo disease.

**Figure 2:**
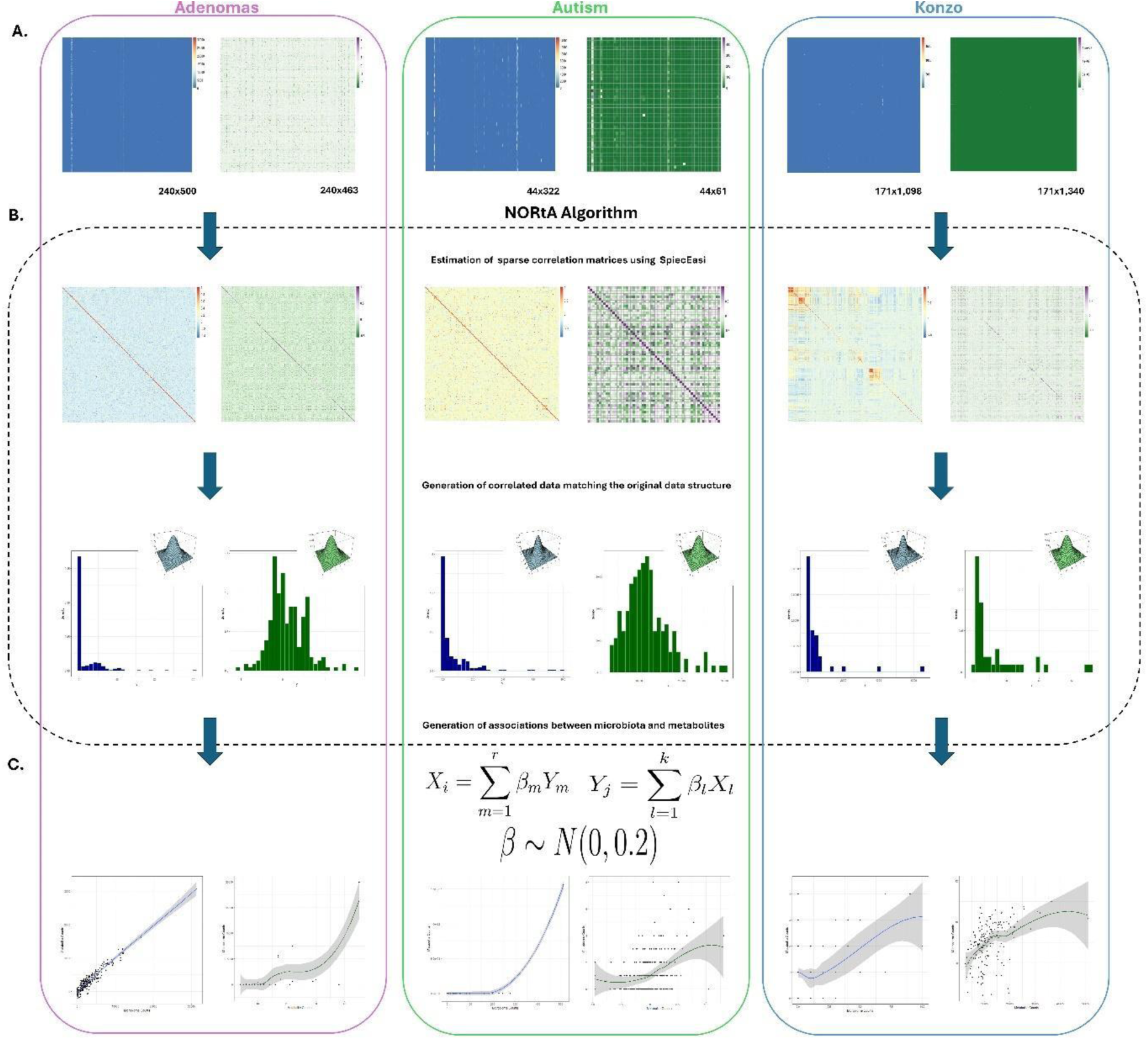
Overview of the simulation setup based on real datasets. (**A.**) Three microbiome-metabolome datasets were selected, each exhibiting different data structures and correlations. We reported the sample size (N) and the number of features (P), as NxP, for each dataset. (**B.**) Realistic datasets were simulated using the “Normal-to-Anything” (NORtA) framework. First, we estimated sparse microbiome and metabolome correlation networks using SpiecEasi ^56^. Second, correlated multivariate Gaussian distributions were generated for both microbiome and metabolome datasets using the correlation structures estimated in the previous step. Third, Gaussian distributions were converted into arbitrary distributions matching the original data structures (**C.**) Associations between species and metabolites were specified mimicking the complex entanglement between the two omics. For each dataset, proportions of associated features vary between 1% and 10%, with association strengths randomly picked from a Gaussian distribution.

### GLOBAL ASSOCIATIONS

A common question in practice for researchers is to find global associations between two omics datasets ^2^. Thus, we compared three multivariate methods detecting associations occurring at a global level between microbiome and metabolome, the Procrustes Analysis ^12^, the Mantel test ^13^, and MMiRKAT ^14^. Since methods have distinct assumptions with respect to the underlying data structures, we provided fair method performance evaluations in two separate settings (See Additional Simulation Settings in the Supplementary Materials). In our main setting, the Mantel test and Procrustes Analysis exhibited comparable results across microbiome data normalizations for controlling the false positive rate in the Adenomas and Konzo scenarios. While in the Autism datasets, the Mantel test considering the Euclidean distance provided the best control of Type-I error rate; the Procrustes analysis and other distance kernels showing inflation patterns (Figure 3A.). Nonetheless, methods highlighted variable powers depending on the scenario, microbiome normalizations and distance kernels considered (Figure 3B.). Indeed, in the Konzo scenario, the alpha transformation could detect more than twice fewer associations compared to the CLR or ILR normalizations, whether the Mantel test or the Procrustes Analysis are considered (Figure 3B.). Additionally, the Canberra distance showed weakest powers across all the three settings, where no clear distinction between Euclidean and Manhattan distances could be observed (Figure 3B.). As an example, in the Adenomas simulation-based scenario, considering the CLR normalization, both the Manhattan and Euclidean distances offered up to 2.72-fold higher powers than the Canberra kernel (Power Manhattan = 68%; Power Euclidean = 67%; Power Canberra = 25%). In an additional scenario with a smaller number of features than individuals, the Mantel test was on average the best method across microbiome data normalizations and distance kernels, showing good control of the Type-I error rate and power (Figures 3C.-3D.). Briefly, when considering the Euclidean distance for the Mantel test, the method was minimally 1.5 times more powerful than MMiRKAT considering the same distance kernel or the Procrustes Analysis (Figure 3D.). This result was consistent across other microbiome normalizations. Our results were confirmed in additional scenarios and considering other types of correlations (Figures S1-S4). Our findings point to different method capabilities to detect global associations depending on the data at-hand. Importantly, our results suggest better performances for the Mantel test regarding both Type-I error rate and power when compared to MMiRKAT or the Procrustes analysis. Particularly, applying the Mantel test using the ILR normalization on microbiome data and the log transformation on metabolites were a robust strategy, offering the best results to detect global associations in a variety of scenarios.

**Figure 3:**
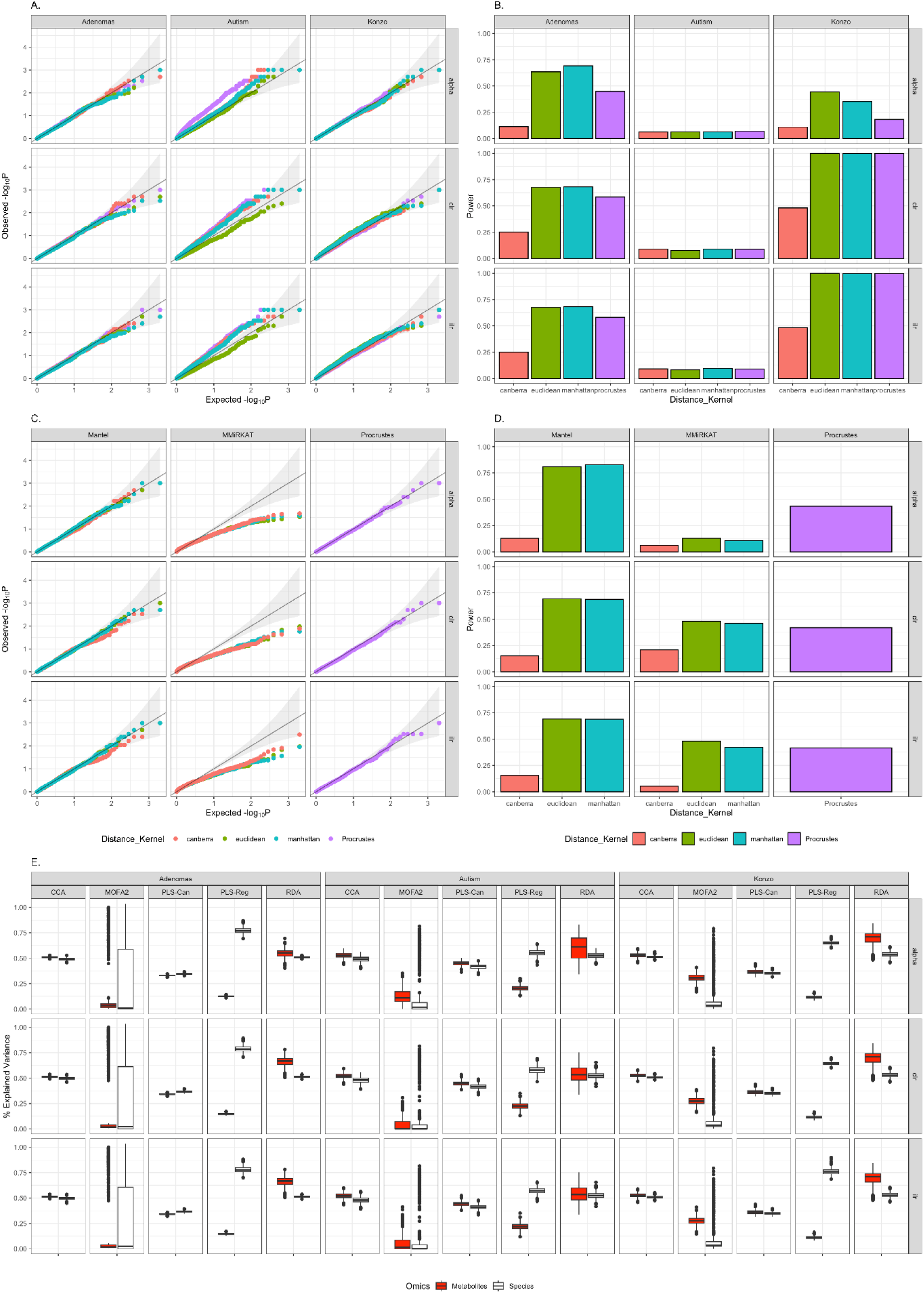
Performance of the multivariate methods for global associations or data summarization. (**A.**) QQ-Plot of the Mantel test and the Procrustes Analysis across microbiome normalizations and distance kernels. For the Mantel test we considered Spearman’s method for computing the global association between the two datasets. P-values for both the Mantel test and Procrustes Analysis were obtained empirically based on 1,000 replicates. (**B.**) Power of the Mantel test and the Procrustes Analysis across microbiome normalizations and distance kernels. For the Mantel test we considered Spearman’s method for computing the global association between the two datasets. P-values for both the Mantel test and Procrustes Analysis were obtained empirically based on 1,000 replicates. P-values <= 0.05 were considered as significant. (**C.**) QQ-Plot of MMiRKAT, the Mantel test and the Procrustes Analysis across microbiome normalizations and distance kernels. Points below the straight line refer to a conservative behavior in the result section. To accommodate MMiRKAT (fewer number of features than sample size), we considered scenarios with a smaller number of features in both omics than number of individuals (See supplementary methods) (**D.**) Power of MMiRKAT, the Mantel test and the Procrustes Analysis across microbiome normalizations and distance kernels. To accommodate MMiRKAT (fewer number of features than sample size), we considered scenarios with a smaller number of features in both omics than number of individuals (See supplementary methods). P-values for both the Mantel test and Procrustes Analysis were obtained empirically based on 1,000 replicates. P-values <= 0.05 were considered as significant. (**E.**) Proportion of explained variance for the data summarization methods across different data structures and normalizations considering the log metabolome. Data summarization methods were compared considering scenarios with a number of features half the number of individuals (See supplementary methods).

### DATA SUMMARIZATION

Instead of measuring one global association, one can be interested in recapitulating information contained within the two datasets through latent factors, accounting for the between-within-correlation ^36^. Thus, we compared Canonical Correlation Analysis (CCA) ^15^, Regression PLS (PLS-Reg) ^16^, Canonical PLS (PLS-Can) ^16^, Redundancy Analysis (RDA) ^17^, and Multi-Omic Factor Analysis (MOFA2) ^18^ across our three main scenarios regarding their capability to summarize the explained variance contained in the two omics (See Methods). Regardless of the considered microbiome data normalization, method’s performances were consistent in the three simulation setups (Figure 2E.). For example, RDA exhibited weak variations of the metabolome explained variances across data normalizations, with standard errors of 0.003, 0.003, and 0.002, for Adenomas, Autism, and Konzo simulation-based scenarios. Impacts of the choice of normalization on result variability for other methods were provided in Figure S5. This result suggests that the normalization used for dealing with compositionality observed in metagenomics data does not affect the capability of methods to capture data variability. Importantly, we observed that no methods are unambiguously the best across data summarizations and simulation settings, where performance seems to depend on the underlying data structure. However, RDA provided consistent performances compared to other methods in most scenarios, showing an average explained variance of 52% depending on the omics of interest (Figure 2E.). Moreover, MOFA2 exhibited the highest variable results spanning from 0% to 100% of explained variances. While surprising, this result suggests that the underlying data structure is critical for the method performance. For regression-based methods, such as PLS-Reg or RDA, the integration directionality impacted the method performances, producing up to 6-fold higher explained variances (Figure 2E.). We confirmed our findings where considering the original metabolome (Figure S6). Our results pointed to RDA as the best trade-off to summarize data variability through latent factors, while PLS-Reg could be exploited when strong assumptions of the effect direction can be assumed. As a whole, our findings suggested that the RDA or PLS-Reg are versatile and robust under scenarios commonly encountered in practice, where the choice of the best method could be driven depending on the research question (See Discussion).

### INDIVIDUAL ASSOCIATIONS FOR COMPOSITIONAL PREDICTORS

Studying the relationship between metabolites and microorganisms may represent an important challenge when accounting for the compositionality induced by microbiome datasets. Indeed, the perfect correlation brought by the compositionality makes the application of standard methods incorrect. This is particularly true when species are incorporated as covariates ^24,29^. We therefore compared four equivalent strategies for studying the global effect of microorganisms on one particular metabolite, HALLA ^37^, the Log-contrast model ^29^, MiRKAT ^14^ and a linear regression on the CLR transformed microbiome (referred to as clr-lm). Globally, under the null hypothesis, across our three scenarios, methods exhibited different behaviors regarding the control of the Type-I error rate, showing an accurate control in the Adenomas simulation-based setting, or being either conservative or slightly liberal in the Autism and Konzo simulation-based scenarios (Figure 4A.). We observed a similar pattern when investigating Pearson’s and Spearman’s correlation with an accurate control or modest inflations of false positives depending on the considered setting (Figure S7). This result suggests a certain discrepancy in method performances to control the Type-I error rate depending on the underlying data structure. At the nominal 5% level, methods highlighted weak to modest powers, ranging from 7% to 35% depending on the method and dataset considered, when these powers were drastically reduced after correcting for multiplicity (Figure 4B., Figure S8). These results are partially explained by the simulation setting where a low signal-to-noise ratio was assumed, mimicking complex microbiome-metabolome relationships. In certain replicate null powers were observed while in others a high percentage of significant associations (Figure S9). Then, comparing models to Pearson’s and Spearman’s pairwise correlations (straight lines in the Figure 4B.), we found no clear advantage of the Log-contrast, HALLA, MiRKAT or clr-lm over standard approaches, when higher powers were mainly due to uncalibrated false positive rates. Interestingly, MiRKAT was robust to microbiome normalization, offering consistent control of the Type-I error rate and power across CLR, ILR, or alpha transformations (Figure 4; Figure S10). Collectively, our results align with a poor calibration of the Type-I error rate in our Konzo and Autism-based scenarios for HALLA, the clr-lm, or log-contrast regression, while MiRKAT exhibited consistent results across the considered scenarios. For all methods, well-behaved qqplots suggesting accurate control of false positives were only observed in the Adenomas simulation-based setting suggesting that the underlying data structure is a key player of method performances. Our evaluation suggested that MiRKAT offers the best false positives-power trade-off in all our simulation settings. The pros and cons of each strategy were further elaborated in the Discussion section.

**Figure 4:**
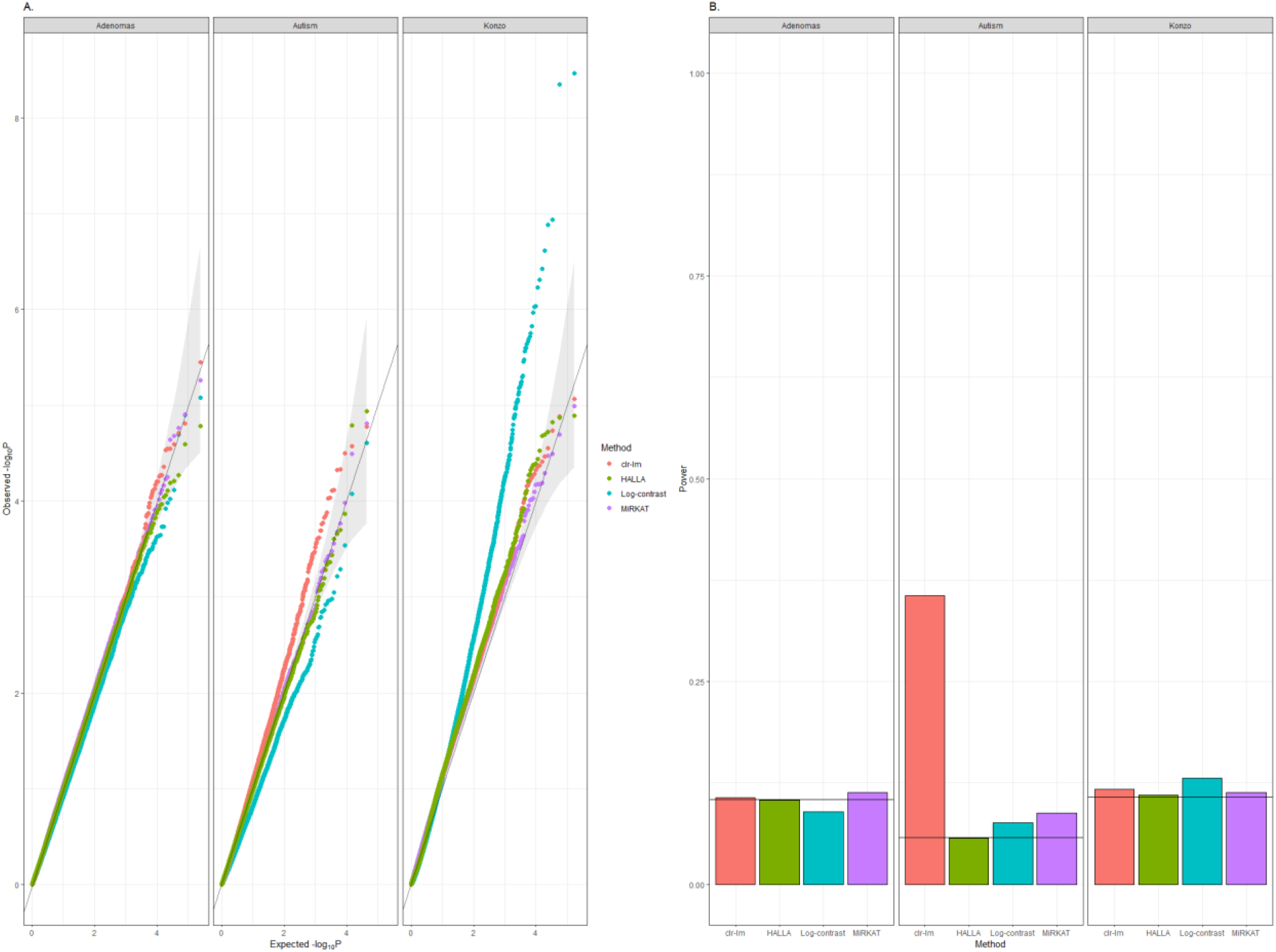
Performance of the individual association methods for compositional predictors. To accommodate long running times due to the number of pairs between species and metabolites we considered scenarios with a number of features half the number of individuals (See supplementary methods) (**A.**) QQplots of the individual association methods across our three simulation settings. (**B.**) Power of the individual association methods across our two main scenarios. P-values <= 0.05 were considered as significant. For the clr-lm method and HALLA, p-values were combined using ACAT ^57^ in order to provide similar comparisons with the log-contrast regression and MiRKAT (See Methods). For MiRKAT, we reported Type-I error rate and power using the ILR transformed microbiome data and the log transformed metabolites, while for HALLA we considered the CLR transformed microbiome and the log metabolome. The straight line represents the background ACAT-combined power using the Spearman’s correlation on the CLR microbiome and the log metabolome. Powers were averaged over 1,000 replicates.

### UNIVARIATE FEATURE-SELECTION FOR COMPOSITIONAL PREDICTORS

Feature selection methods have gained increasing interest from researchers in order to identify a subset of microbiota associated with a variable of interest ^2,30,38^. However, due to the compositionality induced by microbiome data, traditional methods have been shown to lead to incorrect results ^24^. Thus, we compared univariate feature selection methods accounting for compositional predictors, CODA-LASSO ^30^, clr-LASSO ^30^ and clr-MLASSO. First, we evaluated whether methods were able to provide sparse sets of microorganisms across our three scenarios and found that clr-LASSO or CODA-LASSO exhibited smaller sparsity than clr-MLASSO, suggesting that the two methods perform better to keep only a small portion of the original number of species (Figure 5A.). For example, in our Konzo-based simulation setting, we observed that CODA-LASSO offered the sparsest method with on average 0.2% of features selected (sd=0.001) compared to 2% (sd=0.003) and 41% (sd=0.09), for the clr-LASSO and clr-MLASSO, respectively. This result was confirmed in the two other scenarios with different underlying data structures. Then, we assessed how accurate the methods are to find true associations, evaluating methods based on specificity and sensitivity (see Methods). Overall, methods exhibited different patterns of results, with weak to high sensitivities depending on the method and scenario considered (Figure 5A.). As an example, under our three simulation settings, CODA-LASSO showed sensibilities of 0.3%, 10%, and 30%, in the Adenomas-based, Autism-based, and the Konzo-based scenario, respectively, suggesting method performance discrepancies under realistic data structures. We observed comparable results when considering clr-LASSO or clr-MLASSO. Additionally, because the great majority of features were non-associated, methods showed highly specific behaviors. Interestingly, clr-MLASSO due to underlying feature selection process could be highly sensible at the price of lower sparsity or specificity compared to the two other methods. Collectively, our results suggest that univariate feature selection methods must be carefully chosen, depending on the underlying data structure, to accurately select sparse subset of microbiota associated with metabolites (See Discussion).

**Figure 5:**
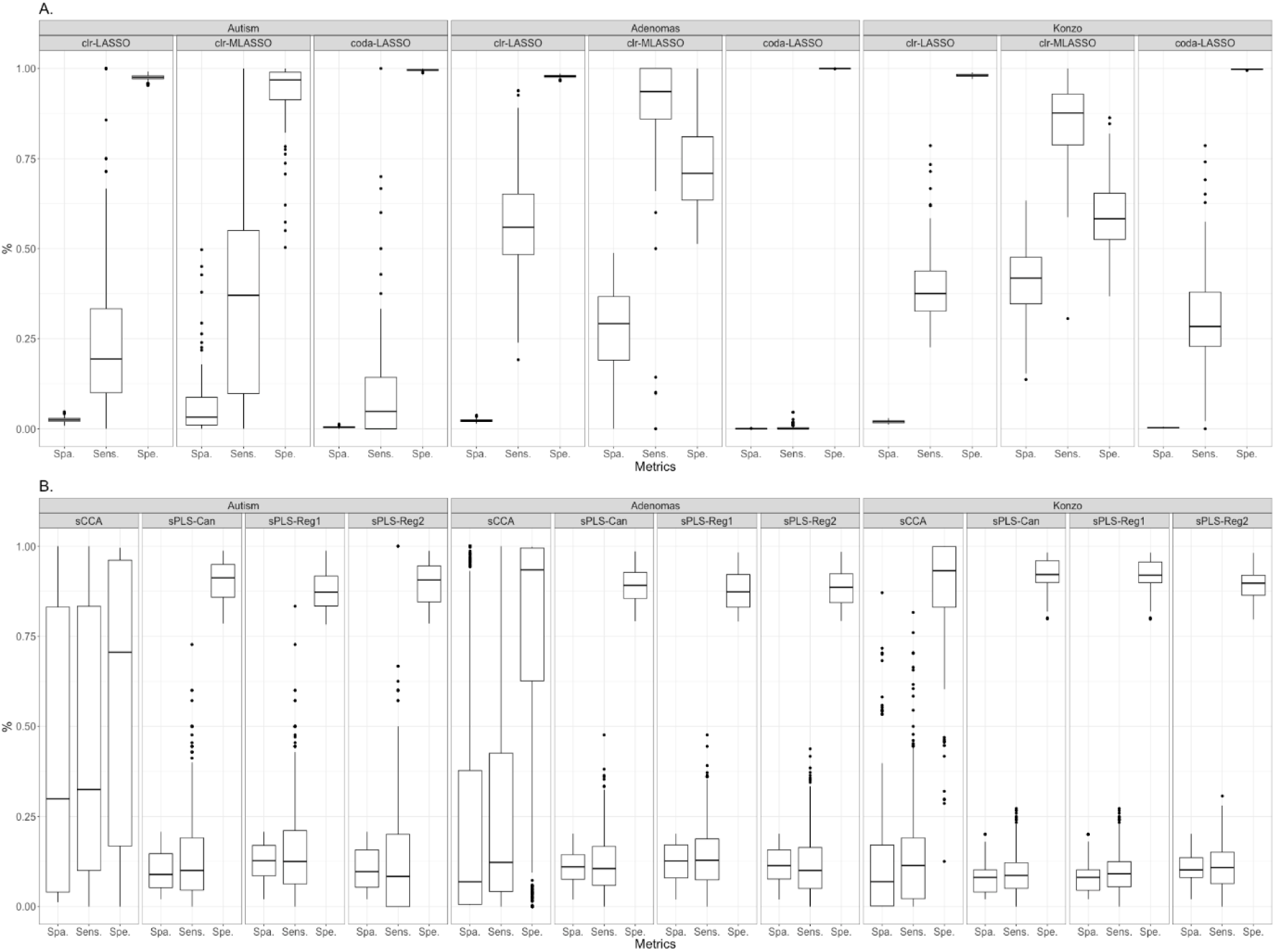
Performance of the feature selection methods for providing sparse and reliable subset of elements across our two scenarios. (**A.**) Performance of univariate feature selection methods considering microorganisms as covariates across our three settings. Metabolites were log transformed before running methods. Performances were calculated on 100 replicates. For CODA-LASSO in the Konzo scenario, we adapted the simulation setting, selecting 300 species and 600 metabolites for accommodating running times of the method (See supplementary methods). (**B.**) Performance of multivariate feature selection methods. Metabolites were log transformed before running methods. sPLS-Reg1 and sPLS-Reg2 correspond to the sPLS-Reg with X=microbiome and X=metabolome, respectively.

### MULTIVARIATE FEATURE-SELECTION

Instead of analyzing each feature independently, exploiting information shared across two omics may represent an interesting avenue to select the most contributive features ^39^. Thus, we compared three methods taking advantage of both intra- and inter-correlation occurring between features of the two datasets, the regression sparse PLS (sPLS-Reg) ^22^, the canonical sparse PLS (sPLS-Can) and the sparse CCA (sCCA) ^21^, respectively. Across our three scenarios, either sPLS-Can or sPLS-Reg provided lower sparsity scores compared to sCCA. Indeed, while sPLS selected on average 12% of the total number of features, sCCA tends to keep either all or no variables, suggesting a poor method’s performance to select features, as confirmed by the patterns of specificity and sensibility (Figure 5B.). This result is higher than the upper bound of true associations assumed by the simulation scenario (10%) suggesting that methods tend to provide a higher proportion of false associations than expected. Further investigations showed that the difference in sparsity may be explained by the distribution of the penalty parameters with severe, high or homogeneous penalization patterns depending on the underlying data structure (Figure S11). Interestingly, both sPLS-Can or sPLS-Reg exhibited the same behavior, showing important levels of specificity and modest sensitivities, while sCCA showed inconsistent performances across our three simulation settings. For example, in the Konzo-based scenario, the two methods offered on average 12% of signals detected as true when the signal is true (sensitivity), with much higher variability for sCCA as demonstrated by the shape of the boxplots in Figure 5B. Finally, when focusing on sPLS-Reg, we found that directionality of integration did not offer discrepancy in results, suggesting robustness to the underlying outcome structure. These results were confirmed when considering metabolites on the original scale (Figure S12). Collectively, our findings align with the conclusion that multivariate feature-selection methods perform moderately to discriminate informative to uninformative features when exploiting the intra-, inter-correlation occurring between omics.

### REAL-DATA APPLICATION

Our systematic evaluation of strategies to jointly analyze microbiome and metabolome data has permitted the identification of the best methods depending on the research question. Thus, we illustrated best approaches through an application on metabolomics and metagenomics data of the Konzo disease ^35^. We presented the exact workflow in the Konzo data analysis section and Figure S13. Firstly, we asked whether there is a different pattern of global association between the two omics in cases and controls and found a stronger relationship in Konzo-affected individuals (Mantel statistic r: 0.4272; Spearman’s permutation p-value: 9.999e-05) than in healthy individuals (Mantel statistic r: 0.2838; Spearman’s permutation p-value: 0.0026997). Then we applied the RDA and found that the two first components explained roughly 26% of metabolome variability across the two conditions, while these proportions remained stable when considering microbiome (Figure 6A.). Moreover, the top-20 most contributing features in each omic on the two first RDA factors highlighted a large panel of associations between microbiota and metabolites with distinct patterns of correlations occurring in affected and unaffected subjects (Figure 6B.; Figure S14). For example, in unaffected individuals, RDA identified *mevalonate* as strongly positively associated with species and *3-hydroxyisobutyrate* exhibiting moderate negative association with microorganisms. These two metabolites have been shown to be linked to inflammatory- or oxidative-based processes potentially involved in Konzo ^40,41^. Consistently, the application of RDA has allowed the detection of 15 *Prevotella* species which have a negative correlation with metabolites in unaffected samples whereas they are positively correlated with metabolites in KONZO affected samples. Moreover, *Bifidobacterium pseudocatenulatum*, *B. adolescentis*, and *B. angulatum* are positively correlated in unaffected samples, whereas *B. pseudocatenulatum*, *B. catenulatum*, *B. longum* are negative in affected patients. A high number of Streptococcus species have a positive correlation uniquely in Konzo affected samples. This result points to distinct patterns of associations between species and metabolites in healthy and affected subjects. Subsequently we used the sPLS regression and identified 30, and 45 metabolites and 235, and 130 microorganisms significantly contributing to the two first components, in cases and controls, respectively (Figure 6C.). Interestingly, of the 130 species kept by the sPLS regression in healthy individuals, 64% were also found in affected individuals, while 46% of metabolites were preserved between the two conditions. Interestingly, both *mevalonate* and *3-hydroxyisobutyrate* have been found to be contributing metabolites in Konzo-affected subjects. To investigate the implication of interactions between metabolites and species in Konzo, we sequentially applied MiRKAT and CODA-LASSO on the subset of unique metabolites found in affected individuals. Consequently, we found that *mevalonate* and *3-hydroxyisobutyrate* were significantly associated with 14 and 16 associated species, respectively both exhibiting a large panel of associations (Figures 6D.-6E.). We identified species which could play a role in Konzo in affecting oxidative response of the metabolism. For example, *Desulfovibrio desulfuricans* was positively associated with *mevalonate*, suggesting that increases in the microbial abundance is associated with an augmentation of metabolite levels. Also, *Clostridioides difficile* exhibited a consistent effect across *mevalonate* and *3-hydroxyisobutyrate,* suggesting common microbial dynamics between the two metabolites. *Clostridioides difficile* has already been reported to have an impact in oxidative stress-related pathways, potentially involved in Konzo ^42^. These associations were missed when applying the clr-lm regression. We validated our findings at a larger scale by a systematic network analysis from the clr-lm regression and CODA-LASSO (Figures 6F.; S15-S17). Our results from metagenomics and metabolomics data from Konzo disease highlight distinct patterns of interactions between microorganisms and metabolites occurring in both affected and unaffected individuals, where different microbial dynamics are involved.

**Figure 6:**
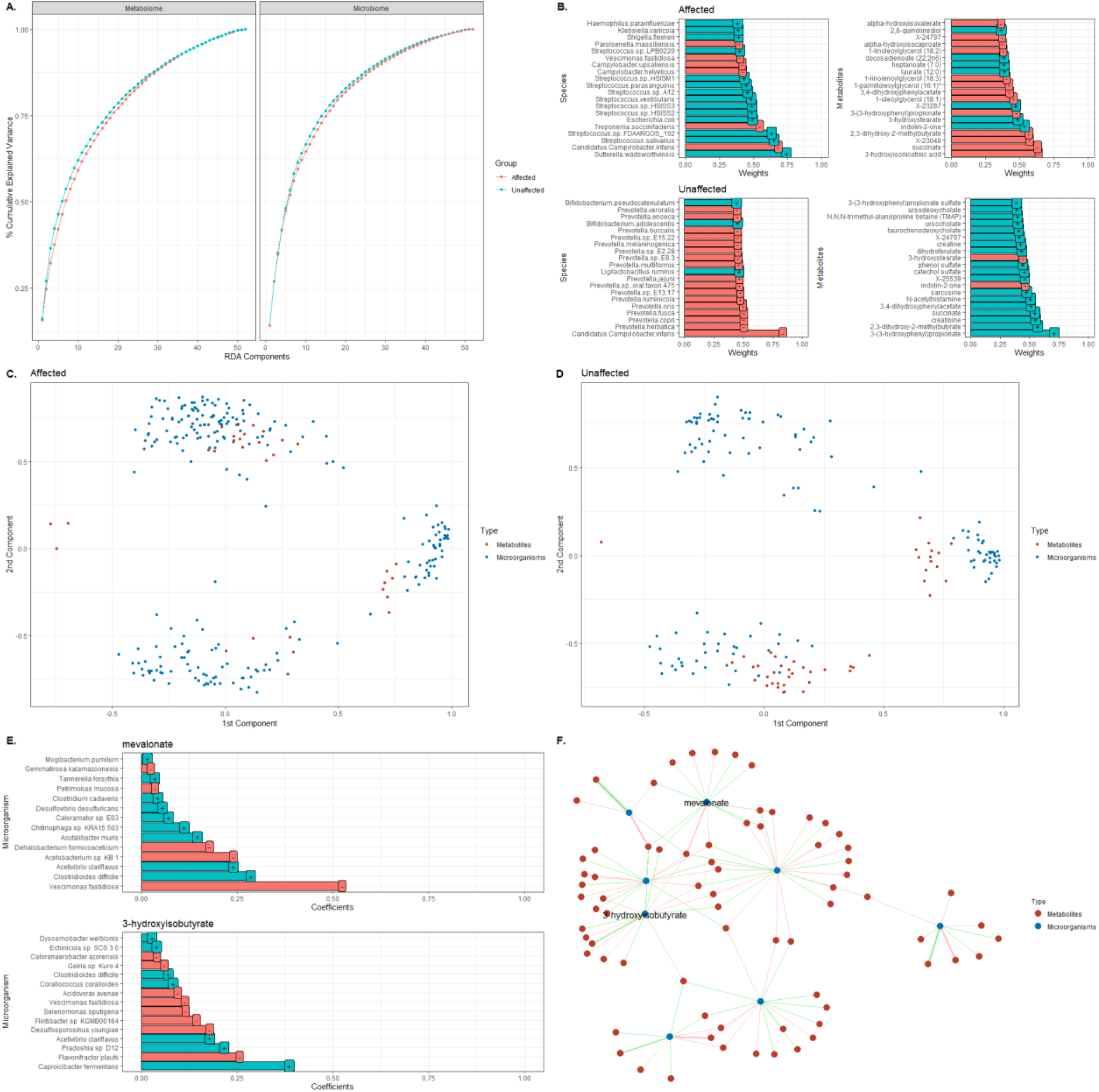
Application of best strategies highlights complementary biological interactions between microorganisms and metabolites in Konzo data. (**A.**) Proportion of cumulative explained variance in Metabolome and Microbiome datasets in both affected and unaffected individuals (**B.**) Top-20 of the most contributing species and metabolites on the first RDA component in healthy and affected samples. Positive correlations were identified by a +, while negative correlations were identified with a – sign. Projection of metabolites (red) and microorganisms (blue) into the 2D regression sPLS space in (**C.**) affected and (**D.**) unaffected individuals. Features with null loadings were removed from the analysis. (**E.**) Coefficients provided by the CODA-LASSO across *mevalonate* and 3*-hydroxyisobutyrate* identified only in Konzo by the regression sPLS. Positive coefficients were identified by a +, while negative coefficients were identified with a – sign (**F.**) Network between *mevalonate* and 3*-hydroxyisobutyrate* and their corresponding associated species found by CODA-LASSO. Positive associations were represented by green edges and negative associations by pink edges.

## Discussion

The integration of microbiome and metabolome datasets within statistical frameworks has become a valuable resource for researchers to comprehensively understand the underlying biological mechanisms involved in diseases. Indeed, recent studies in inflammatory bowel disease ^43^ or cardiometabolic traits ^4^ have highlighted that pathoetiology may result in the disruption in the complex architecture occurring between the two omics. Understanding these interactions represent therefore a critical avenue for unraveling the biology of complex phenotypes. However, currently, there are no standards on how to integrate these two omics together, pushing researchers to constantly waste a lot of time in their decision-making process. Thus, deciding which method fits best for a specific biological question remains a daunting task, critically limiting the result interpretations and replicability. In this paper, we extensively benchmarked nineteen existent integrative methods to disentangle microbiome-metabolome interactions covering most of the researcher aims: *global associations*, *data summarization*, *individual associations*, and *feature selection* (Figure 1). Based on a comprehensive and realistic simulation study and a real data application, we highlighted best methods depending on the research question and data at-hand, providing important insights about statistical good practices (Table 1) and avenues for future methodological developments (Table 2). Furthermore, a user guide and all the scripts used in this work are available in the GitHub repository associated with the paper to facilitate reproducibility and further advancements in the field.

When evaluating global association methods, our results have pointed to important lessons for practitioners. Indeed, across our three realistic simulation scenarios, the Mantel test is the most powerful method to find associations occurring at a global scale compared to MMiRKAT and the Procrustes Analysis. Also, the method exhibits an adequate control of Type-I error rate showing robustness across a variety of scenarios, considering several underlying data structures, normalizations, and distance kernels. This is an appealing feature in practice since choosing the right data transformation or distance metric may represent an important challenge for practitioners. However, by exploiting regression-based frameworks, MMiRKAT, unlike the Mantel test, can adjust for confounding factors making the correction for certain bias induced by individual characteristics, such as age, sex, lifestyle, or even batch effects, possible ^14^. However, the method is unable to deal with scenarios with a larger number of features than individuals, limiting applications in most multi-omics scenarios, as pointed by ^11^. We therefore recommend using Mantel test in most cases or MMiRKAT filtering out features based on a feature selection approach when confounding is expected. The Procrustes Analysis could be exploited to have graphical representations but show no advantages over the Mantel test or MMiRKAT in our benchmark. Importantly, when using the Mantel test, our results suggest that the Canberra distance on metabolome data is the poorest choice for detecting global associations across all our scenarios (Figures 3B.). The application of the Mantel test in our metagenomics-metabolomics use case for Konzo disease has permitted to identify distinct patterns of global associations between microbiome and metabolome occurring in affected and unaffected individuals. Thus, our recommendation here is to apply Euclidean distance on the transformed microbiome data while applying Euclidean or Manhattan distances on metabolites in most cases.

Data reduction is often used by practitioners for summarizing information through a small number of components. Having an efficient method which recapitulates variability across two omics is critical for facilitating subsequent analyses such as visualization or clustering. We considered five different methods exhibiting specific features to summarize omics information and found that in addition to being robust to data normalization, RDA is the most reliable method showing consistency across simulation scenarios and data normalizations (Figure 3E.). Interestingly, our results point to important impacts of directionality when applying regression-based methods, such as PLS-Reg or RDA, with performances drastically increasing depending on whether species or metabolites are considered as the outcome. This result could be explained by the underlying complex structure of microbiome data, strongly impacting the capability of methods to capture data variability. Consistent with this idea, we observed unpredictable performances for MOFA2 with explained variances going from 0% to 100% in all our scenarios (Figure 3E.). Although MOFA2 has been shown to capture complex relationships for gene expression or methylation ^44^, our results align with a distinct pattern of performance for metagenomics and metabolomics data. This behavior is explained by the underlying assumptions of MOFA2, where the use of the method could be restricted to cases where quasi-normality of data is expected, since other types of data require statistical approximations ^18^. We then applied RDA to our Konzo dataset and found distinct microbiota-metabolites associations involved in healthy and affected samples (Figures 6A.-6B.). In unaffected patients the metabolites *3-(4-hydroxyphenyl)lactate* and *5-hydroxyindoleacetate* are identified in positive correlations and may be linked to the fermentation process of the *Bifidobacterium spp*. positively correlated in unaffected samples and negatively correlated in affected samples.

In practice another important question for researchers is to determine the relationship between microbial communities with a variable of interest ^35,45^. However, the underlying compositional structure of microbiome data is an important challenge for model performance. In this paper we have compared four methods accounting for the compositionality of predictors with different strategies, a linear regression applied on the CLR transformed microbiome data, MiRKAT, the log-contrast model, and HALLA. Compared to correlations, these methods have not been shown significantly more powerful (Figure 4B.). As already pointed out in the result section, weak powers are mostly explained by the simulation scenario where we assumed a low signal-to-noise ratio. This behavior could therefore be expected in real-data applications. Thus, we recommend applying univariate methods to only a relevant subset of features, avoiding systematic applications across all metabolite-microbiota pairs. Also, certain univariate methods suffer from inflations of the Type-I error rate in our scenarios (Figure 4A.), suggesting that method performances are impacted by the underlying data structures. For example, the log-contrast model exhibits three different patterns of results across our three scenarios, showing well calibration, conservative, and liberal behaviors in our Adenomas-, Autism-, and Konzo-based scenarios (Figure 4A.). Our general recommendation to the reader is to use MiRKAT, since the method has shown accurate controls of Type-I error rate across microbiome normalizations, permitting correction for confounding factors. This is particularly important since the choice of the right normalization could be difficult. However, microbiome transformations could have critical impacts on result transferability and interpretability, where CLR provides still-correlated synthetic features and ILR or alpha reduce the dimensional space ^30,31^. This result highlights the need for new compositional data transformations, keeping the original number of features while linearly independent (Table 2).

Also, one important contribution of this work is to extensively evaluate feature selection methods. This is particularly crucial for researchers to accurately select metabolites and microorganisms involved in a specific biological context. Our results point to moderate performance of multivariate feature selection methods with inconsistent performances across scenarios and the data transformations considered (Figures 5B.; Figure S12). The best performances are achieved for univariate feature-selection methods for compositional predictors, with CODA-LASSO as a good trade-off between sparsity and classification performances (Figures 5A.), while being sub-compositional coherent. However, all methods provide discrepancies in performance regarding the data structure. For example, CODA-LASSO is strongly affected by the proportion of zeros in microbiome data as suggested by our results in the Adenomas-based scenario. Thus, we recommend in practice to use CODA-LASSO for scenarios with microbial predictors after removing taxa with a high proportion of zeros, to ensure the good performance of the method, whereas sPLS-Reg could be exploited to select features exploiting the inter-omics correlation. We therefore applied both regression sPLS and CODA-LASSO on the Konzo dataset. Regression sPLS has permitted the detection of a distinct set of metabolites and microorganisms in affected and unaffected individuals (Figure 5C.). From the subset of metabolites contributing to Konzo, CODA-LASSO has highlighted different microbial dynamics of effects (Figures 5F.; Figure S17). This result is aligned with the model where microorganisms may be connected to a large set of metabolites. This complex microbiome-metabolome crosstalk has been shown to be associated with diseases ^6^. However, associations found may result in artifact signals since most feature selection methods benchmarked in this paper suffer from lack of sparsity and reliability. This result is aligned with previous reports where authors have shown poor performances of traditional feature selection models ^46^. Indeed, most penalized methods are built upon cross-validation where small perturbations in data may yield drastic changes in results. Similarly to ^46^ extending sparse multivariate or univariate methods to knockoff framework ^47^ or stability selection ^48^ should represent interesting avenues for improving both sparsity and reliability for compositional data ^49^ (Table 2).

Although our work is focused on evaluating methods for inferring associations between metagenomics and metabolomics data, additional work is still required to comprehensively compare methods in a context of prediction. Indeed, predicting metabolome levels from microbiome data is a flourishing research topic with critical implications for clinical applications. Indeed, addressing many challenges, such as integrating and analyzing diverse omics data types, dealing with high-dimensional data, addressing data heterogeneity, and developing robust computational models that capture complex relationships between different molecular layers, could be part of further investigation.

Although our simulation setup can realistically simulate microbiome and metabolome data, our framework is limited to the “Normal-to-Anything” framework. However, as discussed by ^50^, simulating pure compositional data from a Dirichlet distribution induced only a small correlation between features, which is often unrealistic regarding the biology of the microbial communities and metabolites. We therefore promote the correlation, zero-inflation and overdispersion characteristics over a purely compositional structure, with no major impacts expected on the conclusions. Also, methods selected through the benchmark assumed a directionality of effects between species to metabolites, other types of approaches could be considered depending on the biological question. Indeed, in some context, linking metabolome to microbiome using models such as the Dirichlet regression could be achieved^33,34^. We explored the performances of such approaches and found poor results. While we did not elucidate whether the results were explained by the methods themselves or the simulation setting, we decided to omit these approaches in the current version of the benchmark since they are not widely used in practice but could be explored in future works. Our illustration could identify biological species and metabolites involved in Konzo-related processes, our application is however limited to intersection-based analyses between affected and unaffected individuals. We are aware that explicit multi-omics models incorporating the disease information, such as DIABLO ^51^ or MDiNE ^52^ are available. However, future evaluations of differential multi-omics integrative strategies are required to mechanistically link microbiome and metabolome to diseases from a dynamic perspective at large scale (Table 2). We argue this aspect is particularly critical to pinpoint the underlying biological mechanisms hence facilitating precision medicine applications ^53,54^

To summarize, in this paper we provide an extensive benchmark of integrative statistical methods jointly incorporating metagenomics and metabolomics data to infer associations between species and metabolites. We hope this work will represent a wonderful opportunity for the multi-omics community to improve research standards and practices. We anticipate systematic applications of methods identified through this benchmark, as shown by our illustration on the Konzo dataset, on unified metagenomics-metabolomics resources^55^. This aspect is pivotal for identifying common and unique species-metabolites interactions occurring at different scales, required for understanding the underlying biological mechanisms involved in diseases.

## Conclusions

In summary, the present study provides to the multi-omics community one of the largest comprehensive benchmarks of statistical frameworks to jointly integrate metagenomics and metabolomics data. Through an extensive and realistic simulation study, we systematically compared nineteen integrative approaches across most of the research questions encountered in practice. We identified the best methods and illustrated their capability to highlight complementary biological processes involved at different scales with an application to microbiome and metabolome data for Konzo disease. Overall, our study provides a robust and replicable comparative framework of integrative methods. We hope this work will serve as a foundation for setting research standards and the development of new efficient statistical models to mutually analyze metagenomics and metabolomics data.

## Key Points

- We systematically evaluated nineteen statistical methods for jointly analyzing metagenomics and metabolomics data across a wide range of scientific questions, including global associations, data summarization, individual associations and feature selection.
- We provided general guidelines for practitioners for properly analyzing microbiome and metabolome data together, facilitating result interpretation and replicability.
- We illustrated the best methods through an application to metagenomics and metabolomics data in the gut for Konzo disease highlighting complementary biological results.

## Methods

### Simulation setups

Microbiome and Metabolome data were simulated using the “*Normal to Anything*” approach (NORtA), already used for different multi-omics analyses ^50,52,58^. An appealing feature of the NORtA algorithm is to provide a framework capable of simulating data from any marginal distribution while specifying arbitrary correlation structures. Thus, we generated both synthetic microbiome and metabolome data with realistic structures from three different microbiome-metabolome datasets. We selected one high-dimensional metagenomics-metabolomics dataset from a recent study in Konzo disease ^35^ exhibiting correlated and over-dispersed microbiome counts, while metabolomics data highlight correlated count data. Under this scenario microbiome data were assumed to follow a negative binomial distribution, and metabolome a Poisson distribution. Then, we chose an intermediate-size microbiome-metabolome dataset in Adenomas ^59^ where species are correlated and zero-inflated counts with a general zero-inflated negative binomial underlying structures, when metabolome were correlated log-normally data. Finally, we picked one small metagenomics-metabolomics from autism spectrum disorder ^60^, where species have correlated zero-inflated negative binomial structures and metabolites a Poisson distribution. We estimated sparse correlations networks of species or metabolites using SpiecEasi as already used in a variety of contexts ^56^. We simulated correlated multivariate Gaussian distributions where the correlation matrices were estimated in the previous step. Normal distributions were converted into any correlated arbitrary distributions using the copula space matching the original data structure. These steps were summarized in Figure 1 of the main manuscript. Several data transformations for both microbiome and metabolome were evaluated across our scenarios to account for data structure. Briefly, we evaluated the impact of three microbiome normalizations (CLR, ILR, and alpha transformations) on method performances. Data normalizations were summarized across methods and simulation scenarios in Table S1. To evaluate the Type-I error control, we independently generated two datasets under the null hypothesis of no association between microorganisms and metabolites. Under the alternative hypothesis, we varied both the number of associations between microorganisms and metabolites and the strength of associations, mimicking microbiome-metabolome complex interdependence. Methods were compared under three realistic scenarios, where both the numbers of features (species or metabolites), the sample size, and the underlying data structures varied. Technical details on the simulation and additional scenarios were provided in the supplementary. Under all scenarios we simulated 1,000 replicates. Simulation setup was summarized in Figure 2.

### Statistical Analyses

Let’s assume X and Y, a matrix of microbiome and metabolome, collected on the same set of samples, of size n x p and n x q, where n is the number of samples, p the number of microbiota and q the numbers of metabolites, respectively. Xij represents the jth microorganism in the ith sample, with j = 1,2,.., p, while Yik is the kth metabolite in the ith sample, where k=1,2,…,q. For the sake of simplicity we considered the case where p=q.

### Global Associations

In this paper we refer to global association methods, the statistical approaches providing global associations between microbiome and metabolome data (Figure 2). As already presented in Deek et al., 2024, we considered three general methods, the Mantel test ^13^, MMiRKAT ^14^, and the Procrustes Analysis ^12^, considering a variety of data transformations and distance kernels.

The Mantel test ^13^ is a statistical framework measuring global correlation between two datasets measured on the same set of samples. Traditionally, the Mantel test is applied on distance or dissimilarity matrices. Here we considered three different distance kernels applied on the original and log-transformed metabolome dataset, Euclidean, Canberra and Manhattan distances. Also, we applied the Euclidean distance on a normalized microbiome matrix, since this projection leads to more natural interpretations ^24^ (Table S1). The Mantel test was applied considering either Pearson’s or Spearman’s correlation, to account for linear and rank-based associations. P-values were obtained empirically based on permutations using 1,000 replicates. The Mantel test was performed using the *vegan* R package.

MMiRKAT is the multivariate extension of MiRKAT providing global association between a distance-transformed microbiome dataset (referred to as kernel function) and a low dimensional continuous multivariate phenotype ^14^. Briefly, the model regresses the multivariate outcome in terms of the non-parametric kernel transformed microbiome up to a gaussian error term. Consistent with distance kernels used in the Mantel test, we considered Euclidean, Canberra and Manhattan distances applied on the transformed microbiome data as kernel-based transformations, while the entire original or log transformed metabolome matrix was considered as the outcome (Table S1). Significance testing is performed by computing kernel specific p-values using the exact davies method. MMiRKAT was applied using the `MiRKAT` R package.

Procrustes Analysis is a method facilitating high-dimensional visualisation of two or more datasets. Traditionally, the method linearly translates, scales and rotates principal components minimizing the Euclidean distance between the two omics (referred to as Procrustes superimposition). Since the two datasets have been superimposed inter-omics distances can be computed for each row, where smaller distances refer to higher adequation between the two omics. Formal testing is achieved by empirically computing the p-value using 999 permutation samples, comparing the observed sum of squared deviation (square of the difference between distances of the same sample) and the resampled ones.

As introduced in the *simulation setups* subsection, when applying the global association methods, we normalized microbiome data using CLR, ILR, and alpha transformations, while metabolome data were considered on the original and log scale.

### Data Summarization

In this benchmark, we considered 4 distinct data summarization methods, encompassing CCA, PLS, RDA, and MOFA2. Briefly, all these methods seek to summarize data information through latent factors.

CCA initially proposed by ^15^ summarizes the relationship between two datasets by finding linear combinations of the two matrices maximizing the correlation. CCA was performed using the `CCA` R package.

Unlike CCA, PLS seeks for linear combinations maximizing the covariance between the two datasets ^16^. Also, in PLS directionality of effect of one matrix on the other can be taken into account, leading to two general forms of PLS, regression and canonical, respectively ^17^. Thus, canonical PLS and regression PLS were applied with the `mixOmics` R package.

Moreover, RDA is a two-step procedure, combining multivariate linear regression and PCA ^17^. In the first step, a multivariate linear regression is fitted between each element of the matrix of responses and the matrix of predictors. Then a PCA is applied on the matrix of predicted values. RDA was performed using the `vegan` R package.

Finally, MOFA2 is an unsupervised multi-omics framework able to untangle sources of variability shared by different omics ^18^. MOFA2 is a Bayesian probabilistic model able to find latent factors linking two omics by putting priors on model parameters. We applied MOFA2 using the related R package `MOFA2` with default parameters.

Except for MOFA2 where the best number of latent factors were chosen by the model, we kept all the components corresponding to the minimal number of features observed in one dataset.

### Individual associations

When individual relationships are of interest, we consider different regression models taking into account the compositionality induced by microbiome data as predictors.

Indeed, for microbiota that are explanatory variables, we fitted four different models, a log-linear regression on the CLR transformed microbiome, a log-contrast model ^29^, MiRKAT^14^, and HALLA ^37^.

Formally the log-linear model of the CLR transformed microbiome (referred to as clr-lm in the Result section) is given by:

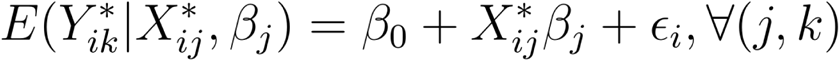

where *Y*\* is the log transformed metabolome matrix and *X*\* the CLR transformed microbiome data. Although the compositionality in the microbiome data is taken into account using the CLR transformation, the previous model is not robust to the subset of microorganisms, not preserving the sub-compositionality feature of microbiome data. Thus, the log-contrast model by imposing a zero-sum constraint on regression coefficient preserves the scale invariance property needed to ensure the sub-compositionality characteristic of microbiome data ^29^. Formally, the model is given by:

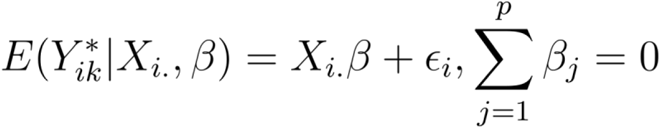

Under the log-contrast framework, following ^29^ we applied the global significance F-test in order to determine whether there is an association between at least one microorganism and a given metabolite. The log-contrast model was performed using the *Compositional* R package. Aligned with the idea of global association, MiRKAT is a statistical framework exploiting semi-parametric kernel machine regression framework in order to summarize microbiome relationships ^14^. One major feature of MiRKAT compared to other approaches is permitting the use of several distance kernels at the same time. This is particularly appealing since it is often unclear in practice which kernel is the more suitable. In our context, we considered Euclidean, Canberra and Manhattan distances either on original or transformed microbiome data, while considering the original or log transformed metabolome as outcome. MiRKAT was applied with the `MiRKAT` R package. Finally, we evaluated HALLA, a hierarchical All-against-All statistical framework ^37^. The model was initially developed to incorporate both homogeneous and heterogeneous datasets and permits the clustering of features of different omics adequately controlling the False Discovery Rate. HALLA was applied through the corresponding *`halla`* python package with default parameters, considering the CLR-transformed and original microbiome and the log-transformed and original metabolome.

### Feature Selection: Univariate

Adapted from ^30^ we considered two different models accounting for compositional predictors, when fitting models with metabolites as outcomes. Firstly, we considered the CLR-LASSO, performing the CLR transformation on microbiome data before fitting a univariate or multivariate LASSO log-linear regression ^20^. We referred to LASSO and MLASSO in the Results section. Formally for a metabolite k, the LASSO log-linear model is given by:

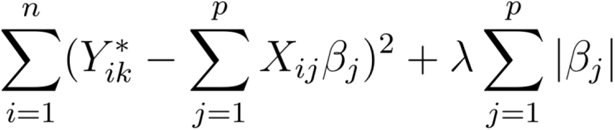

with *Y*\* is the log transformed metabolome matrix and *X*\* the CLR transformed microbiome data. Best penalty parameters λ were chosen using a 10-fold cross-validation through a 10 step grid-search from 0.01 to 1. LASSO or MLASSO models were fitted using the `glmnet` R package.

Then, consistently with the log-contrast model, we applied the coda-LASSO considering a log-linear response of the metabolome level. Briefly, the coda-LASSO is a penalized log-contrast model, permitting to select only the most contributive features, with a zero-sum constraint on regression coefficients, ensuring scale invariance, a property needed for compositional data. The model considered in the coda-LASSO framework is a direct extension of the model initially proposed by ^61^. This latter fits a two-stage model on all possible log-ratios between each pair of microbiota, leading to sparse solutions. The R package `coda4microbiome` with the default parameters were used when applying coda-LASSO.

### Feature Selection: Multivariate

Sparse Canonical Correlation Analysis (sCCA) ^21^ and sparse Partial Least Squares (sPLS) ^22^ are two penalized extensions of CCA and PLS permitting to summarize data information through latent factors while proceeding to feature selection.

For sCCA we used L1 penalty on the two datasets, only keeping features contributing on the two first components. Best penalty parameters were found using 25 permutation-based samples considering a 0.1-step grid search from 0.01 to 1. sCCA were performed using the `PMA` R package.

Consistently, canonical and regression sPLS were tuned using a 10-fold cross validation, considering a 5 step grid search ranging from 5 to 25 in our low dimensional setting and from 10 to 50 in our high dimensional scenario. We maximally kept two components in order to select the most contributive features. sPLS were applied using the `mixOmics` R package. For both sCCA and sPLS, features on the two first components with non-null loadings were considered as informative variables hence were kept to compute the performance metrics.

#### Data and Distance Kernel Transformation

Most methods used in practice need either a normalization step or a distance-based transformation in order to be applied properly on compositional or over-dispersed data. Thus, we considered in our main analyses three data normalizations for microbiome and one data transformation for metabolome data. The choice of data normalization depends on the research objective. We referred to specific data and distance kernel transformations for each family of methods in the subsection *Statistical analysis*.

To consider the compositionality of microbiome data while keeping the original number of features, we considered the centered log-ratio transformation (CLR) ^62^ applied on the original count data. This normalization was considered across all the different considered methods. The CLR transformation computes the log ratio of each microbiota count on the geometric mean for a given individual. Formally, the CLR transformation is given by:

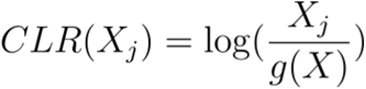

where *g*(*X*) is the geometric mean over all the microorganisms for one sample. This transformation projects the simplex onto a D compositional subspace under a zero-sum constraint ^31,63^. By keeping the original number of features the CLR transformation is a one-one transformation, facilitating result interpretation which is an appealing feature in practice. We therefore considered the CLR transformation as the reference normalization when individual associations or feature selection are of interest. However, the CLR transformation does not ensure independence between features and sub-compositionality coherence. This latter represents a major limitation for distance-based methods due to singular covariance matrices. Thus, when distance between features is of interest, we consider the isometric log- ratio (ILR) ^32^ and alpha transformation ^31^. Intuitively, these two transformations project the original D-dimensional space into an independent D-1 quasi-orthogonal space, the main difference laying into the transformation used. The ILR transformation projects the original data onto a Euclidean space. Formally:

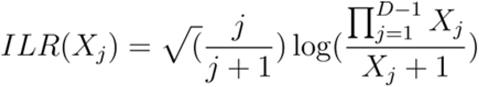

While the alpha transformation is a Box-Cox type transformation, where the transformed data follow a multivariate distribution after a suitable alpha-transformation ^31^.

This facilitates the use of traditional multivariate methods. We therefore considered the ILR and alpha transformations when evaluating global associations, and data summarization methods, since the correspondence with the original features does not really matter. Moreover, since the metabolome data have been shown to be log-normally distributed we applied a natural log transform on the original count data ^26^.

Also, we applied different distance kernel transformations before performing some global association or individual association analyses, highlighting different patterns of relationships occurring among features. Briefly, we considered Euclidean, Canberra and Manhattan distances on metabolome matrices of original and log transformed counts, while considering the Euclidean distance on original and transformed microbiome data. Interestingly, as presented by ^24^, the Euclidean distance applied on CLR transformed data corresponds to the Aitchison distance. This latter has been shown superior to the Bray-Curtis dissimilarity, representing a true linear relationship, while more stable to data subsetting or aggregating ^24^, and will be considered as our reference method here. All data and distance kernel transformations depending on the method used were summarized in Table S1.

#### Performance Metrics

Since all the methods considered in this benchmark exploit different statistical concepts, the outputs cannot be directly compared. Consequently, we opted for several performance metrics depending on the research question.

Indeed, for global and individual association methods, we systematically evaluated model performance through Type-I error control and power, since the considered methods are frequentist frameworks. Briefly, Type-I error control assesses whether a method provides a good control of false positives at a given significance threshold. In other words, under the null hypothesis of no association, at a significance threshold equals to 0.05 we maximally expect 5% of false positives for a method that performs well. A value above the expected rate is considered as liberal, while a value below conservative. Type-I error control was evaluated using the quantile-quantile plot of the −log10 of p-values. Similarly, the power is the capability of a method to detect a significant signal (at a given significance threshold) when we know that there is an association. In practice, researchers want methods maximizing the power while accurately controlling the Type-I error.

Data summarization methods were compared based on the proportion of the explained variance. We refer to explained variance, the amount of data variability kept by latent factors built by methods.

Moreover, inspired from ^46^ when univariate and multivariate feature selection methods were evaluated, we considered sparsity and reliability as primary performance metrics. For univariate methods sparsity corresponds to the total number of relevant associations found by the method (here with coefficients different from zero), while reliability is the capability of a method to accurately discriminate true from false associations between two features. However, we adapted both sparsity and reliability calculation when considering multivariate feature selection methods. Indeed, sparsity was computed by the total number of nonzero coefficients on the total number of features while reliability was adapted to capture the model performance to keep true contributive variables within the two datasets. Reliability was evaluated using the specificity and sensitivity. In practice, researchers are looking for sparse methods with methods discriminating both the true associations from false associations (high sensitivity) and true non-associations from false non-associations (high specificity). Performance metrics depending on the considered method were summarized in Figure 1. Technical details on the performance metric calculation and adaptations were provided in the next subsections.

#### Explained Variance

In order to provide fair comparisons between data summarization methods we extended the redundancy index initially proposed by ^64^ for Canonical Correlation Analysis (CCA) ^15^ to Partial Least Squares (PLS) ^16^. Following ^64^, the redundancy index is the proportion of explained variance kept in the latent factors. See ^64^ for more details on the redundancy index calculation. Although exhibiting structural differences, i.e., CCA maximizes the correlation while PLS the covariance, we argue that extending the redundancy index to PLS provides a good proxy of the data variability contained in the components.

#### Specificity/Sensitivity

When comparing feature selection methods, we used the specificity and sensitivity to determine the classification performance for the models considered. Briefly, the specificity is a metric of prediction performance computed based on the ratio between the number of true negatives and the total number of negative elements. Equivalently, the sensitivity is computed based on the ratio between the number of true positives and the total number of positive elements. From a standard confusion matrix:

| Predicted/Ground Truth | Non-Associated | Associated |
| --- | --- | --- |
| Non-Associated | #True negatives | #False negatives |
| Associated | #False positives | #True Positives |

The specificity is obtained by: #True negatives/(#True negatives + #False positives) and the sensitivity by: #True positives/(#True positives + #False negatives).

For univariate feature selection methods, the specificity and sensitivity are obtained using the true set of associations between taxa and metabolites, i.e, with coefficients different from zero, while for multivariate feature selection methods, metrics are computed with the true set of associated features, i.e, with loadings different from zero.

#### Sparsity

When evaluating the feature selection methods, we compared methods using sparsity. Briefly the sparsity is the method ability to provide the as smaller as possible set of features, either with coefficients or loadings different from zero, depending on the considered method.

Formally the sparsity is given by:

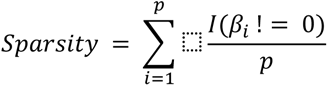

where *β* is either the coefficient for an association from a univariate feature selection method or corresponds to a loading in a multivariate feature selection. p is the total number of possible associations/features.

#### P-value combinations

In order to provide fair comparisons across our individual association methods with compositional predictors, we combined p-values using the Aggregated Cauchy-based test (ACAT) ^57^ when CLR-lm, HALLA, and correlations were considered. Indeed, for a large number of microbiota and metabolites, applying univariate methods can lead to p x q possible correlations, limiting the statistical power due to multiplicity. Similarly to the log-contrast model or MiRKAT, in practice one can be interested in having the global association between one metabolite and several microorganisms. Thus, in order to provide a powerful method controlling the Type-I error rate well, we combined p-values for all microorganisms in a given metabolite using ACAT ^57^, resulting from p p-values. We argue that this approach may result in more detected signals, since the multiplicity burden is drastically reduced. Briefly, ACAT is a method combining p-values through a Cauchy distribution.

Formally for one metabolite, the aggregated p-values across the p microbiota can be approximated by:

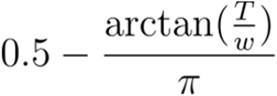

where 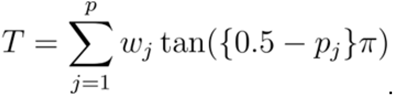

One important feature of ACAT compared to other aggregation methods, such as Fisher’s method, is that the method can efficiently control the Type-I error rate even in presence of correlated p-values, while maintaining good power ^57^. Also, the method does not require any resampling step, facilitating its application to large datasets.

#### Konzo data analysis workflow

Stool samples collected from individuals from study populations in Masi-Manimba (n = 65) and Kahemba (n = 106) regions of the Democratic Republic of the Congo were used for metagenomics and metabolomics assessment, where a proportion of the cohort is affected with Konzo. Shotgun metagenomics sequencing was performed on DNA extracted from ~250mg of stool with the goal of generating ~50 million reads per sample. Data was analyzed following a similar methodology as described previously using Kraken2 and Bracken for taxonomic classifications ^35^. Additionally, stool was analyzed by the company Metabolon, harnessing their large in-house repository of rigorously tested and validated metabolites that are used as reference, to detect metabolites present in the samples. Analysis was performed on the 1,098 microorganisms and 1,340 metabolites across the 171 individuals unconditionally of the disease status. Microbiome data at the genus level were normalized using the CLR transformation while metabolome data were log transformed. The workflow was as follows 1) global association, 2) data summarization 3) univariate and multivariate feature selection and 4) individual associations. Moreover, we considered microorganisms as explanatory variables and the microorganisms as outcomes. For global associations, since the number of features exceeds the number of individuals, we performed the Mantel. Then, we applied RDA in order to detect the most contributing microorganisms and metabolites on the first components. Following the same methodology as presented in the Method section, we extracted the core microorganisms and metabolites using the regression sPLS, keeping only the features with nonzero loadings on the two first components. We finally applied the clr-lm regression and CODA-LASSO in order to highlight contributions of microorganisms on metabolites. We summarized the workflow in Figure S13.

## Ethics approval and consent to participate

Not applicable

## Consent for publication

Not applicable

## Availability of data and materials

Codes to reproduce the analyses are available at: https://github.com/lmangnier/Benchmark_Integration_Metagenomics_Metabolomics. Data to run the code are available at: https://doi.org/10.6084/m9.figshare.25234915. The simulated data are produced using the simulate_data.R script available in the same Github repository. R 4.2.2 is required to reproduce results from the paper. Data for metagenomics and metabolomics for Autism and Adenomas were available at https://github.com/borenstein-lab/microbiome-metabolome-curated-data. The metagenomics and metabolomics data for Konzo disease are available upon request from Matthew S. Bramble.

## Authors’ contributions

LM designed, conducted, performed the data analysis, and wrote the manuscript. AB and MM performed the data analysis. AM, AB, MPSB and NV wrote the manuscript. AB, MPSB, MSB, and AD revised the manuscript. All authors read and approved the final version of the manuscript.

**Loïc Mangnier** is a research fellow in the Arnaud Droit Lab affiliated to the Laval University

**Margaux Mariaz** is an intern in Biostatistics in the Arnaud Droit Lab affiliated to the Laval University

**Neerja Vashist** is a postdoctoral fellow in the Arboleda Lab, in the Department of Pathology and Laboratory Medicine at UCLA

**Antoine Bodein** is a research fellow in the Arnaud Droit Lab affiliated to the Laval University

**Marie-Pier Scott Boyer** is a research fellow in the Arnaud Droit Lab affiliated to the Laval University

**Alban Mathieu** is a research fellow in the Arnaud Droit Lab affiliated to the Laval University

**Matthew Bramble** is an associated professor at The George Washington University School of Medicine and Health Sciences

**Arnaud Droit** is a full professor in the Department of molecular medicine at Laval University

## Acknowledgments

We would like to thank members of the Arnaud Droit Lab, particularly Louis-Maël Gueguen, Thomas Jeanne, and Tania Cuppens for their insightful comments on the manuscript.

