## Supplementary Material for "Decoding the Microbiome-Metabolome Nexus: A Systematic Benchmark of Integrative Strategies"

### Supplementary Materials

**Figures** **1**
Global Associations 1
Data Summarization 4
Univariate Associations 6
Multivariate Feature Selection 10
Real Data Application 12
**Table** **17**
**Methods** **19**
Simulation setting 19
Additional simulation settings 21
**References** **23**

### Figures

#### Global Associations

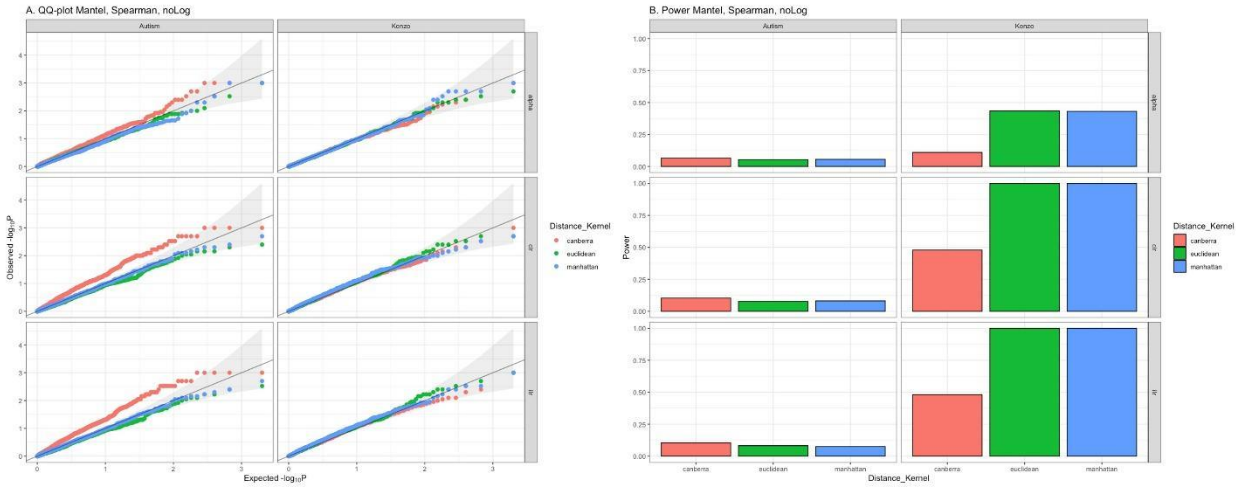

**Figure S1**: QQplots and power for the Mantel test considering Spearman's correlation on the original metabolome data across microbiome normalizations. P-values were obtained empirically based on 1,000 replicates. P-values <= 0.05 were considered as significant.

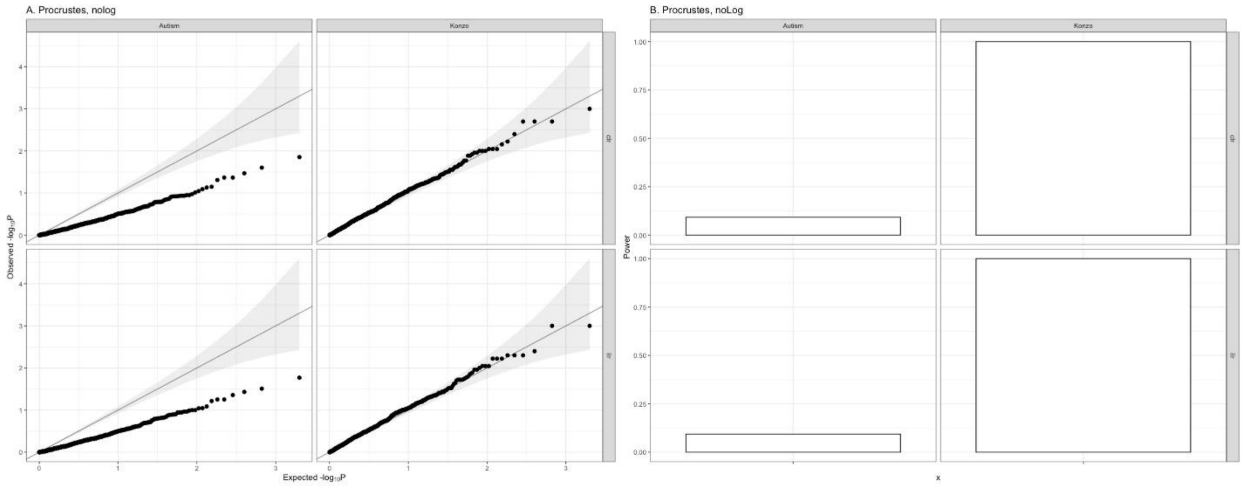

**Figure S2**: QQplots and power for the Procrustes Analysis on the original metabolome data across microbiome normalizations. P-values were obtained empirically based on 1,000 replicates. P-values <= 0.05 were considered as significant.

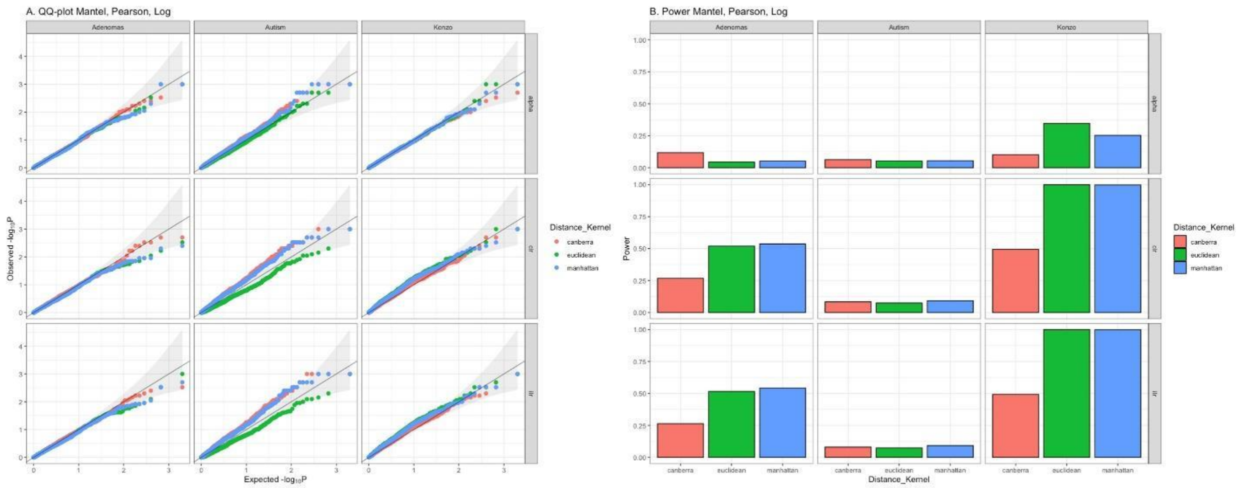

**Figure S3**: QQplots and power for the Mantel test considering Pearson's correlation on the log metabolome data across microbiome normalizations. P-values <= 0.05 were considered as significant.

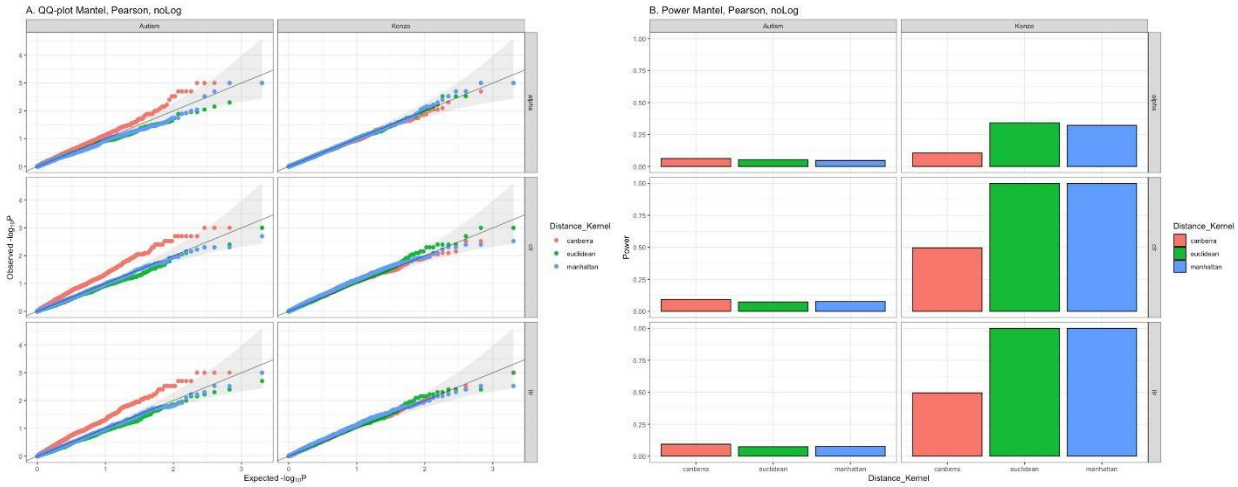

**Figure S4**: QQplots and power for the Mantel test considering Pearson's correlation on the original metabolome data across microbiome normalizations. P-values <= 0.05 were considered as significant.

### Data Summarization

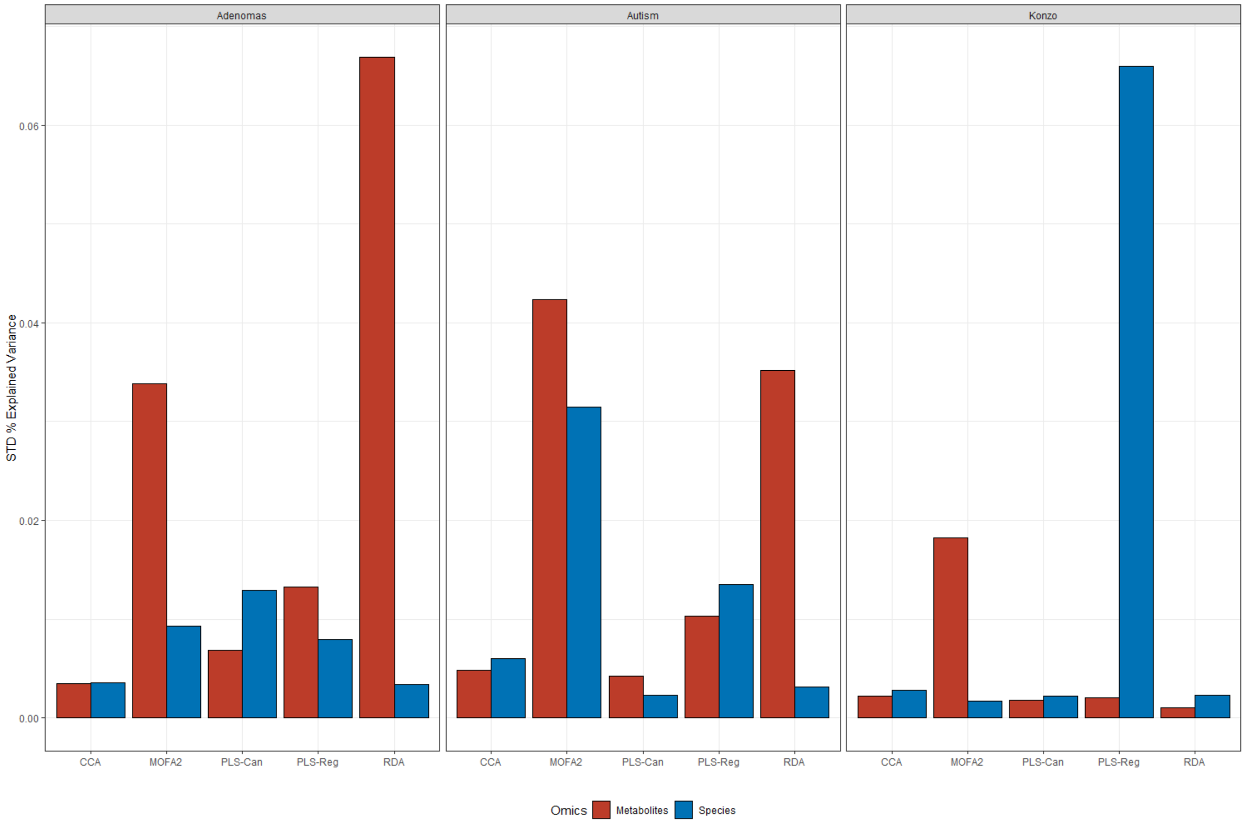

**Figure S5**: Variability performances for data-summarization methods across scenarios considering the log metabolome

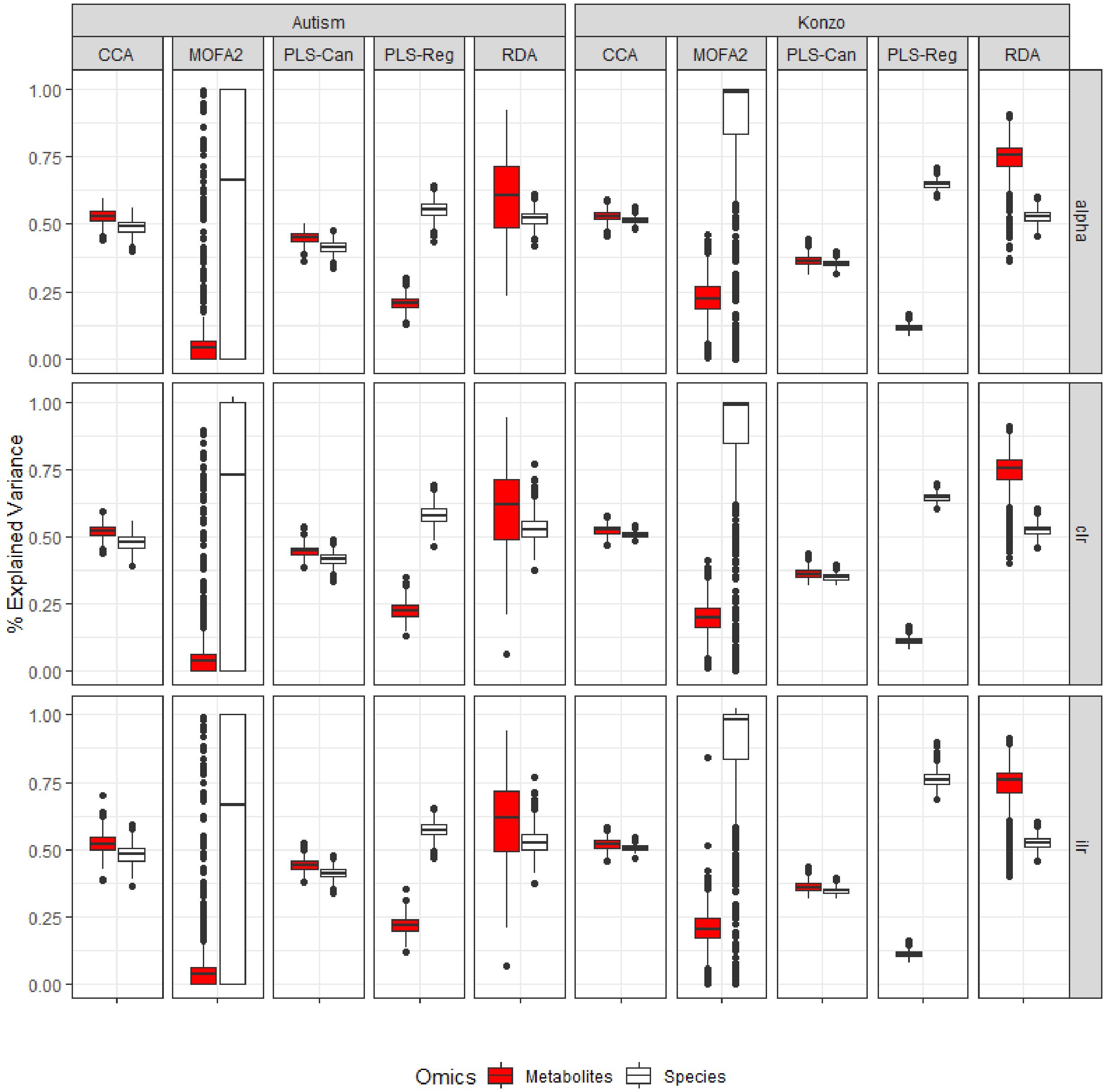

**Figure S6**: Proportion of explained variance for the data summarization methods across different data structures and normalizations considering the original metabolome.

#### Univariate Associations

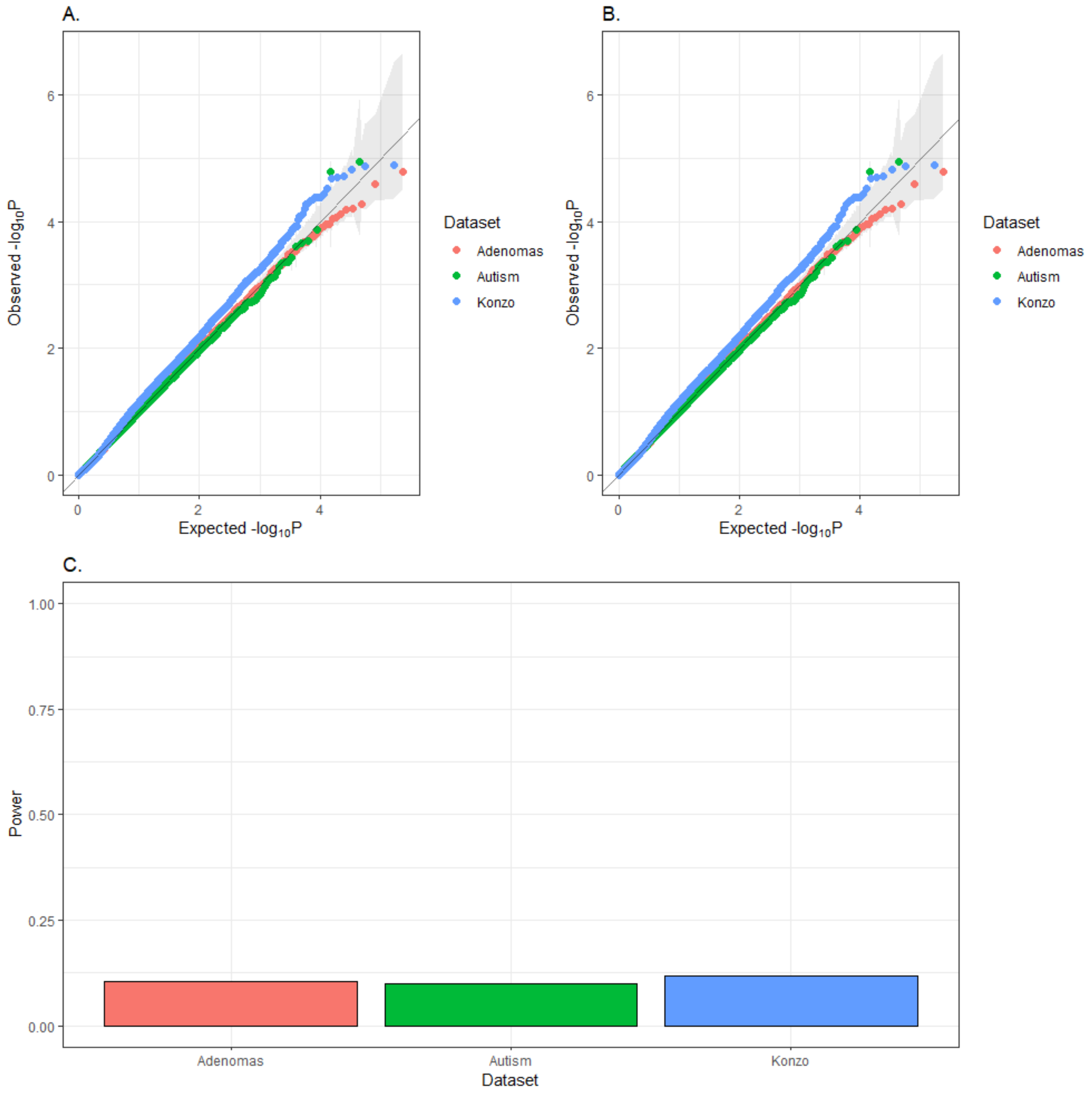

**Figure S7**: (**A.**) QQ plot of the Spearman's correlation on the CLR transformed microbiome and the log metabolome in our three simulation settings. (**B.**) QQ plot of the Pearson's correlation on the CLR transformed microbiome and the log metabolome in our three simulation settings. (**C.**) Power of the Pearson's correlation on the CLR transformed microbiome and the log metabolome in our three simulation settings. P-values <= 0.05 were considered as significant. Powers were averaged over 1,000 replicates

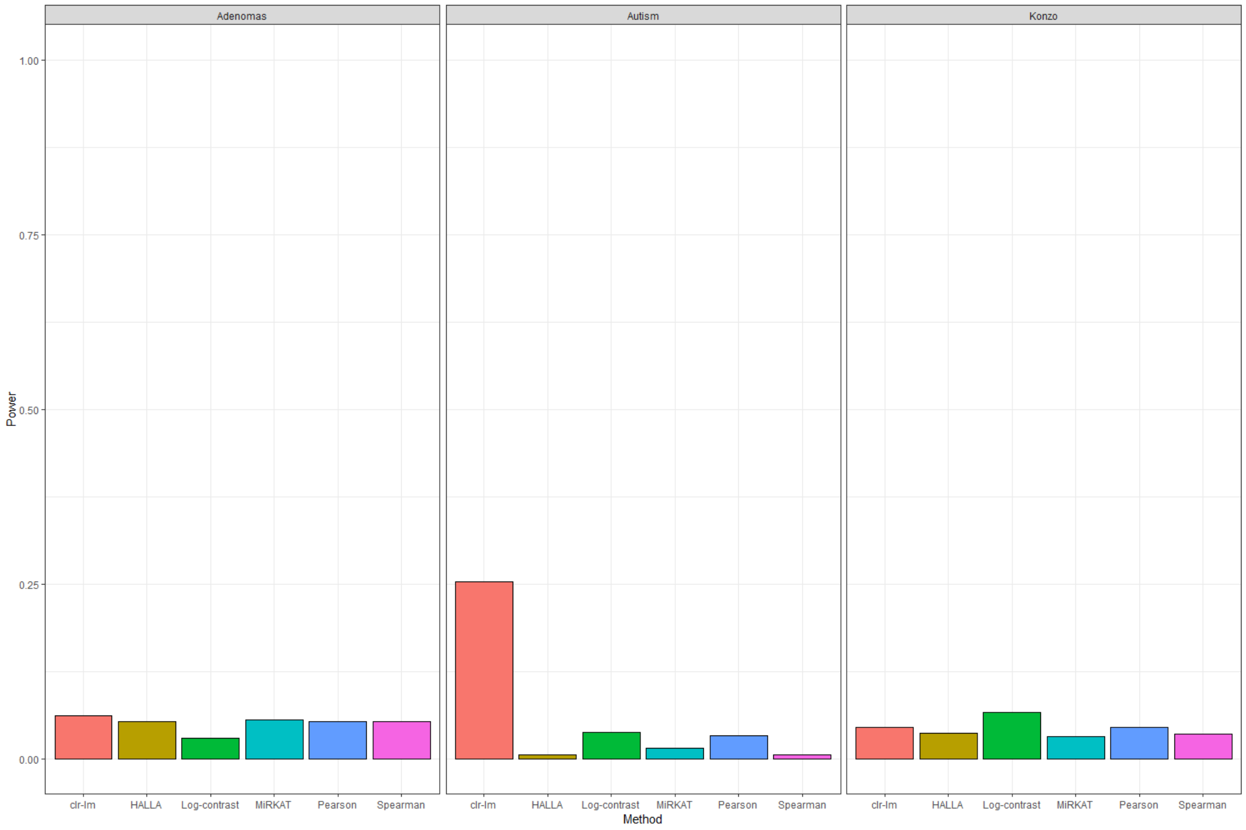

**Figure S8**: Power of univariate methods after controlling for multiplicity computing the FDR across replicates. FDR <= 0.05 were considered as significant. MiRKAT was applied on the ILR transformed microbiome, while HALLA on the CLR normalized microbiome. We considered the log metabolome for all methods. Powers were averaged over 1,000 replicates.

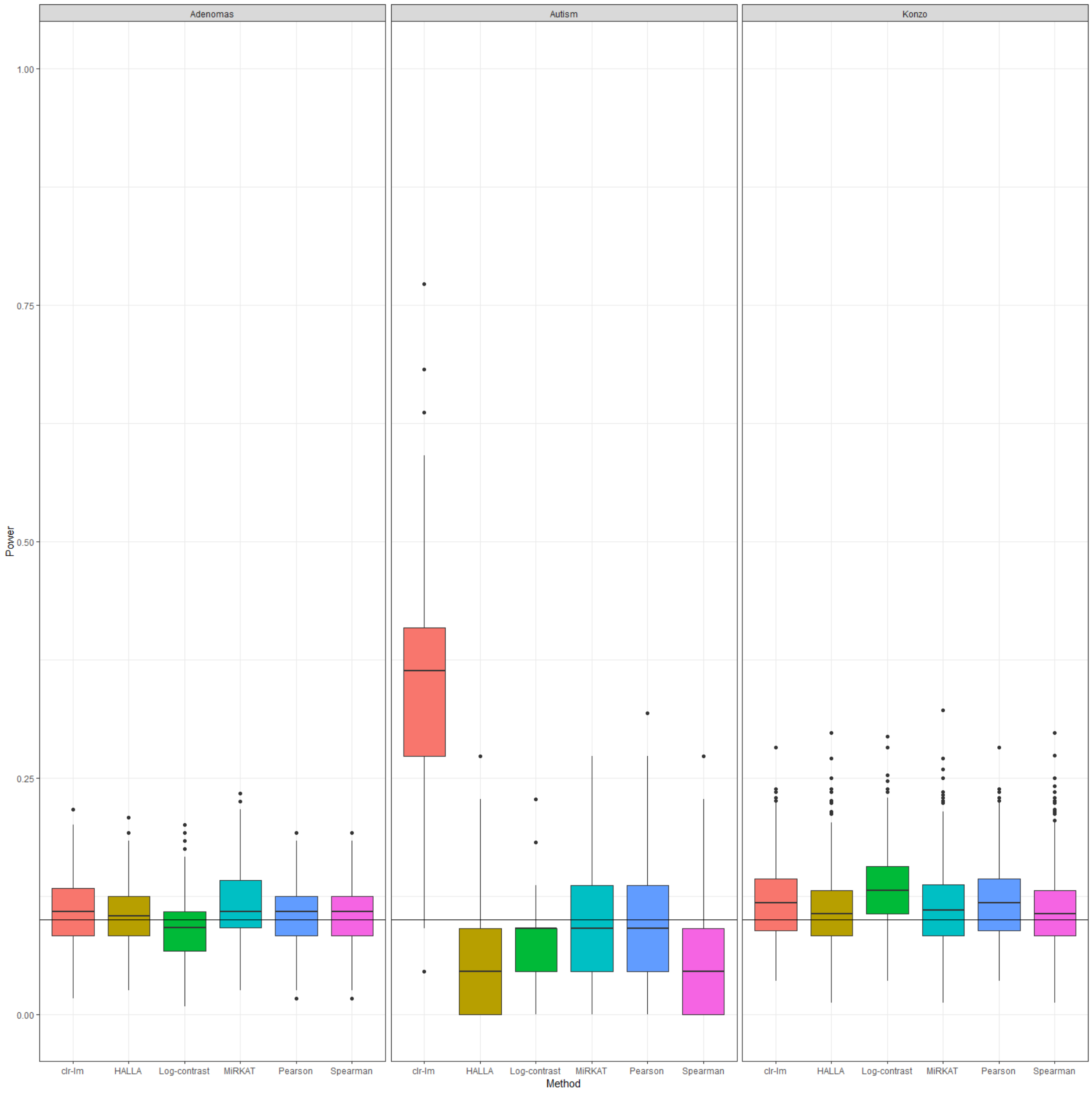

**Figure S9**: Boxplots of powers across simulation settings considering univariate association methods. Straight horizontal line corresponds to the maximal percentage of associations. P-values <= 0.05 were considered as significant.

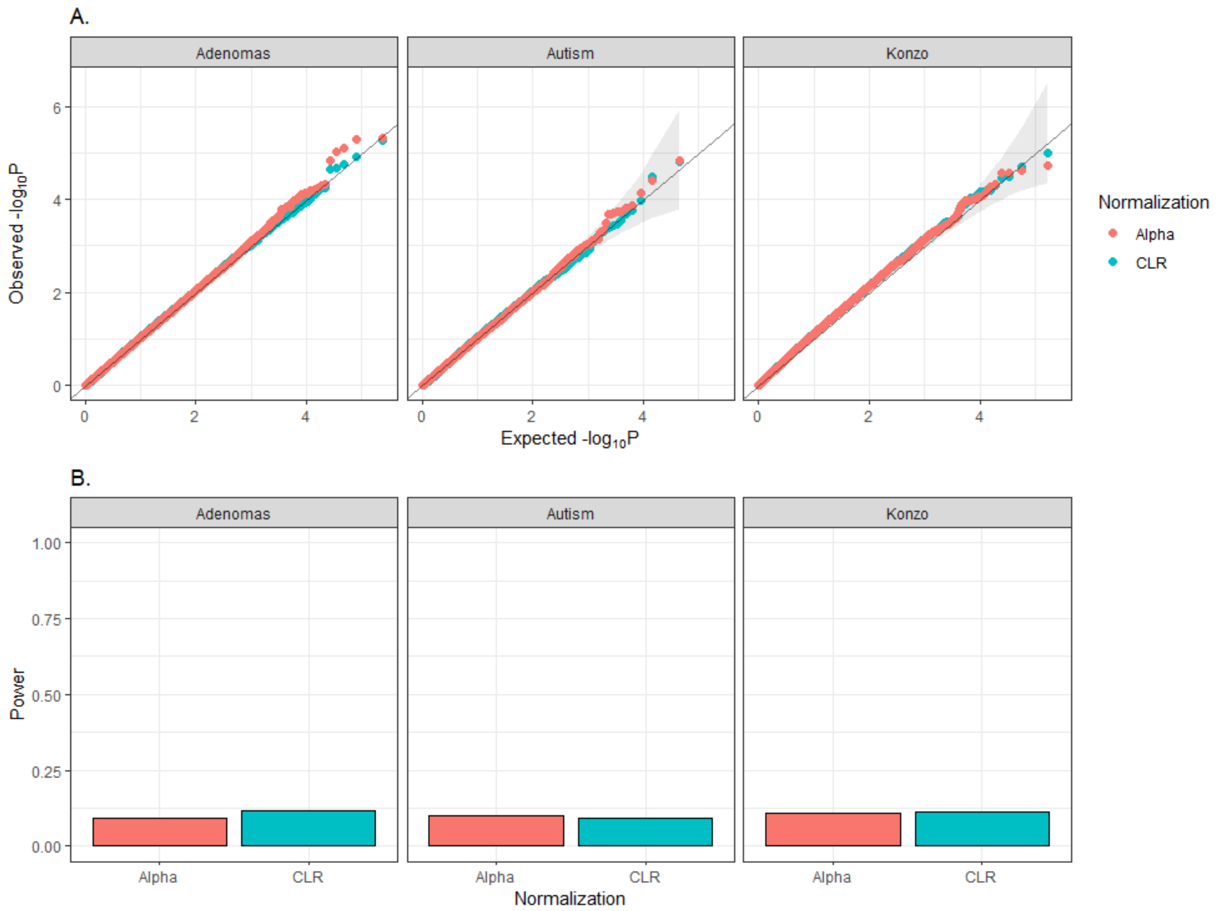

**Figure S10**: (**A.**) QQplots and (**B.**) power for MiRKAT across different microbiome normalizations. P-values <= 0.05 were considered as significant. Powers were averaged over 1,000 replicates.

### Multivariate Feature Selection

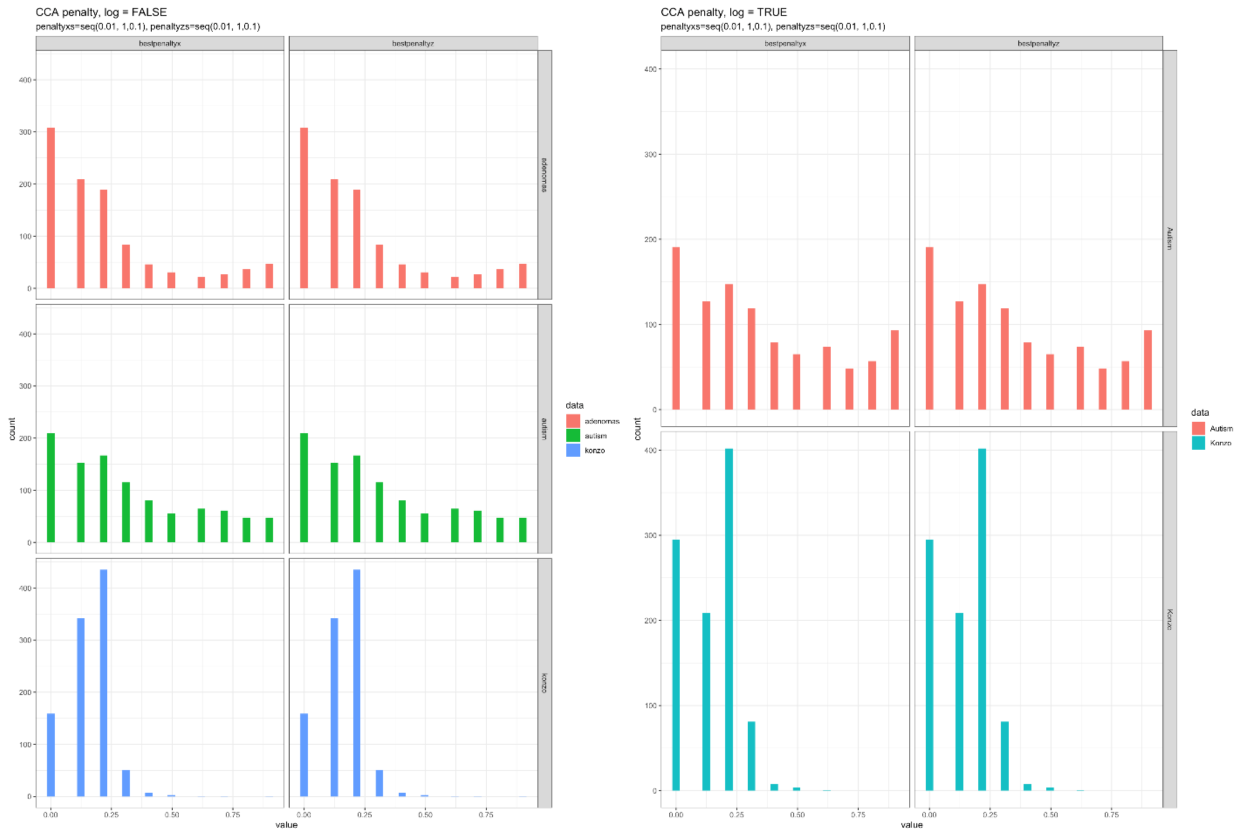

**Figure S11**: Distribution of the best penalty parameters across simulations chosen by sCCA in microbiome (x) and metabolome (y) data considering the original metabolome (left) and log-transformed metabolome (right). Adenomas-based scenario was not considered when log-transformed the metabolome, since metabolites were assumed to be log-normal under this setting. In our simulations, sCCA provided equal penalty parameters for the two omics explaining the same distributions.

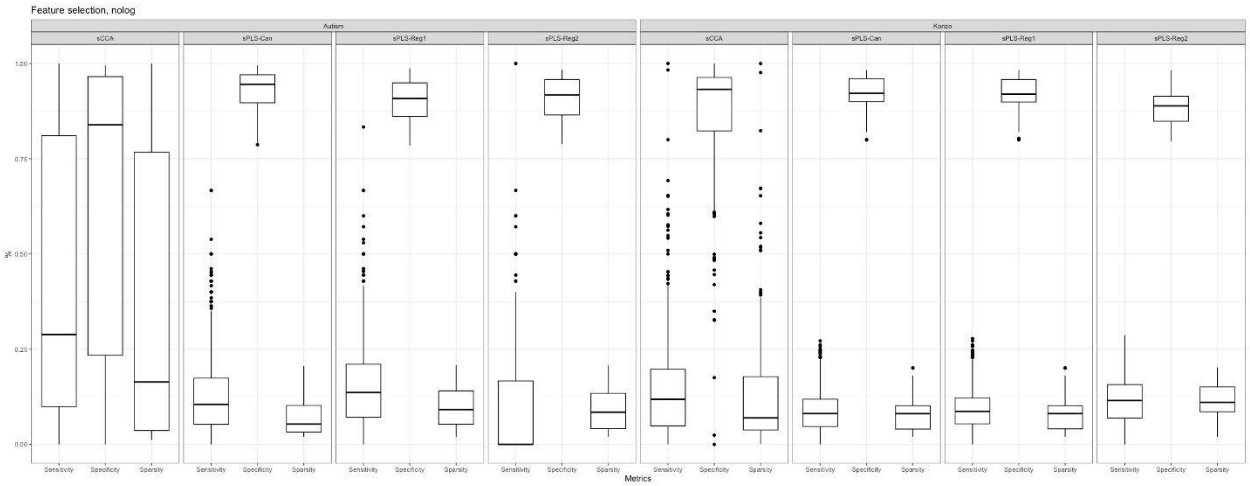

**Figure S12**: Performance metrics for multivariate feature selection methods when considering the original metabolome data. Adenomas-based scenario was not considered here, since metabolome data have already been reported in Figure 5 of the main manuscript. sPLS-Reg1 refers to the sPLS-Reg with metabolome as outcome, while sPLS-Reg2 refers to the sPLS-Reg with microbiome as outcome.

### Real Data Application

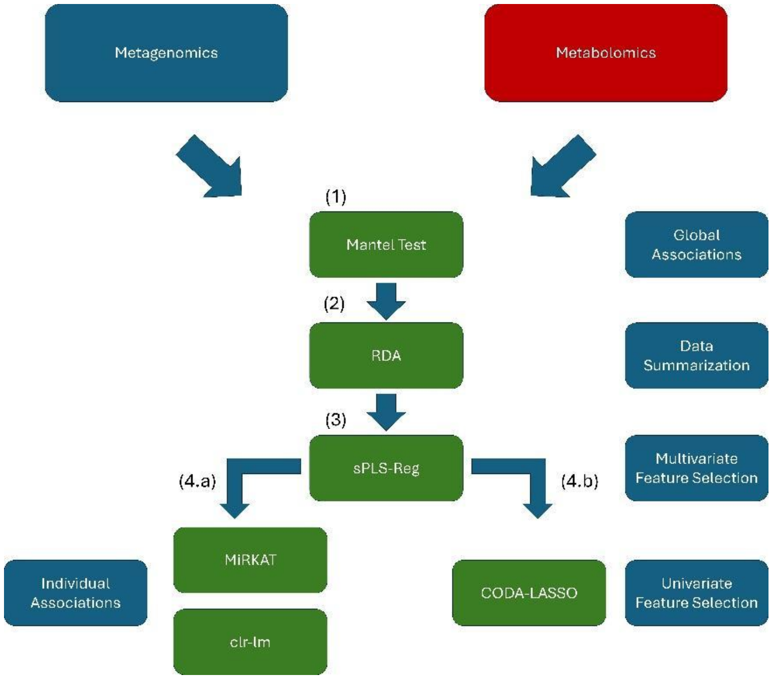

**Figure S13:** Workflow for the Konzo data application. As described in the main manuscript we applied (1) the Mantel test to uncover associations between metagenomics and metabolomics occurring at the global scale (2) RDA to identify features contributing the most to the two first components (3) sPLS-Reg to subset the core species and metabolites exploiting the inter-omics correlation (4.a) MiRKAT and clr-lm and (4.b) CODA-LASSO to identify species associated with metabolites. This workflow was applied in both affected and unaffected individuals.

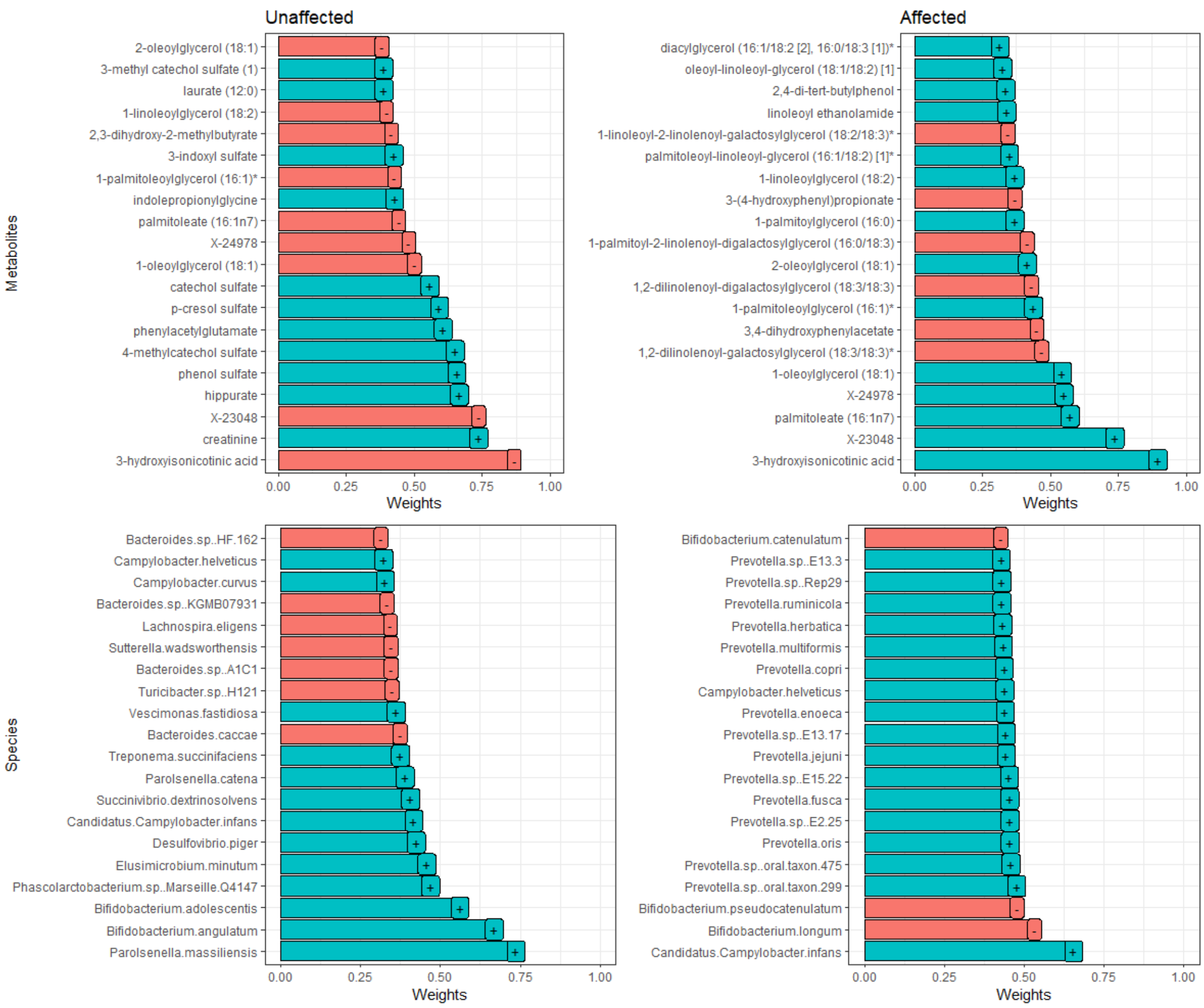

**Figure S14**: RDA weights for the top-20 metabolites and species on the second RDA component in both affected and unaffected individuals

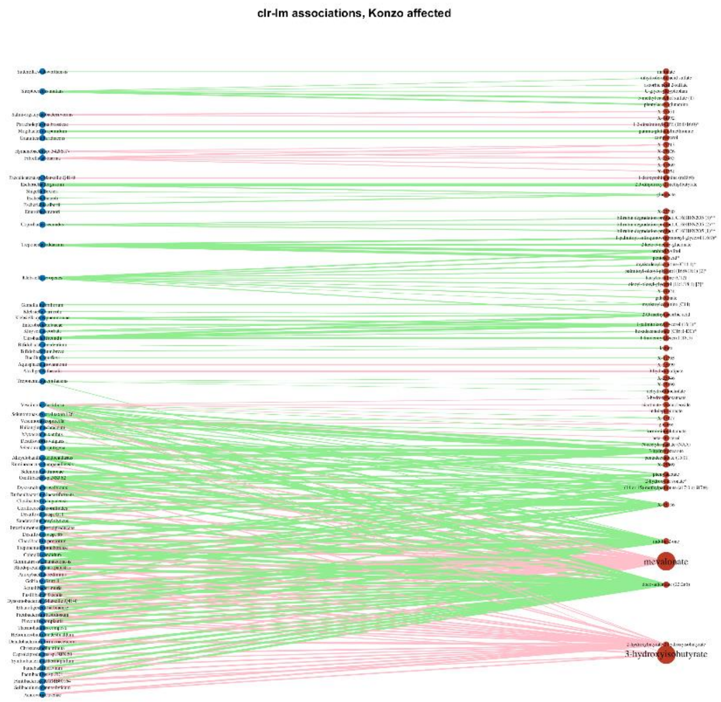

**Figure S15**: Networks of interactions between metabolites (red) and species (blue) provided by the clr-lm regression in affected samples. Only metabolites with an ACAT-Combined p-values were reported. Pink edges represent negative associations while green edges represent positive associations.

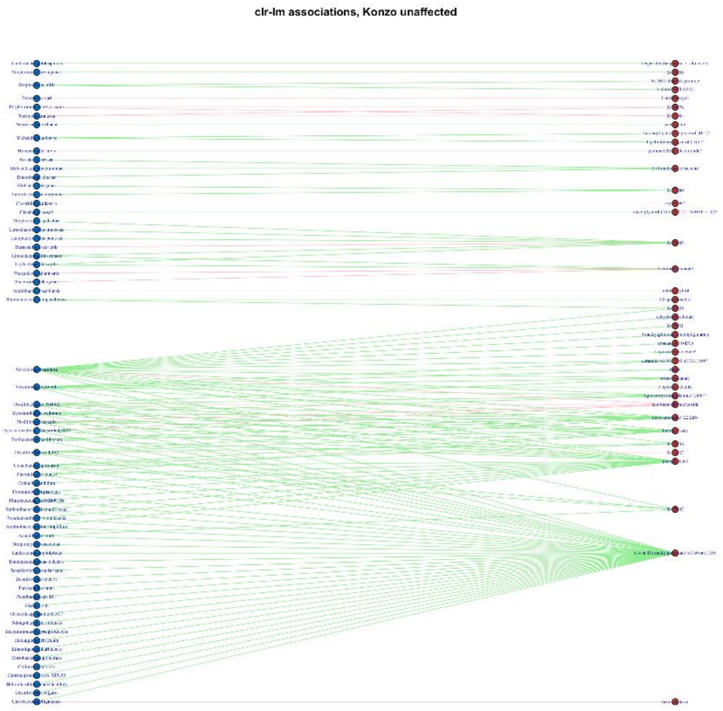

**Figure S16**: Networks of interactions between metabolites (red) and species (blue) provided by the clr-lm regression in unaffected samples. Only metabolites with an ACAT-Combined p-values were reported. Pink edges represent negative associations while green edges represent positive associations.

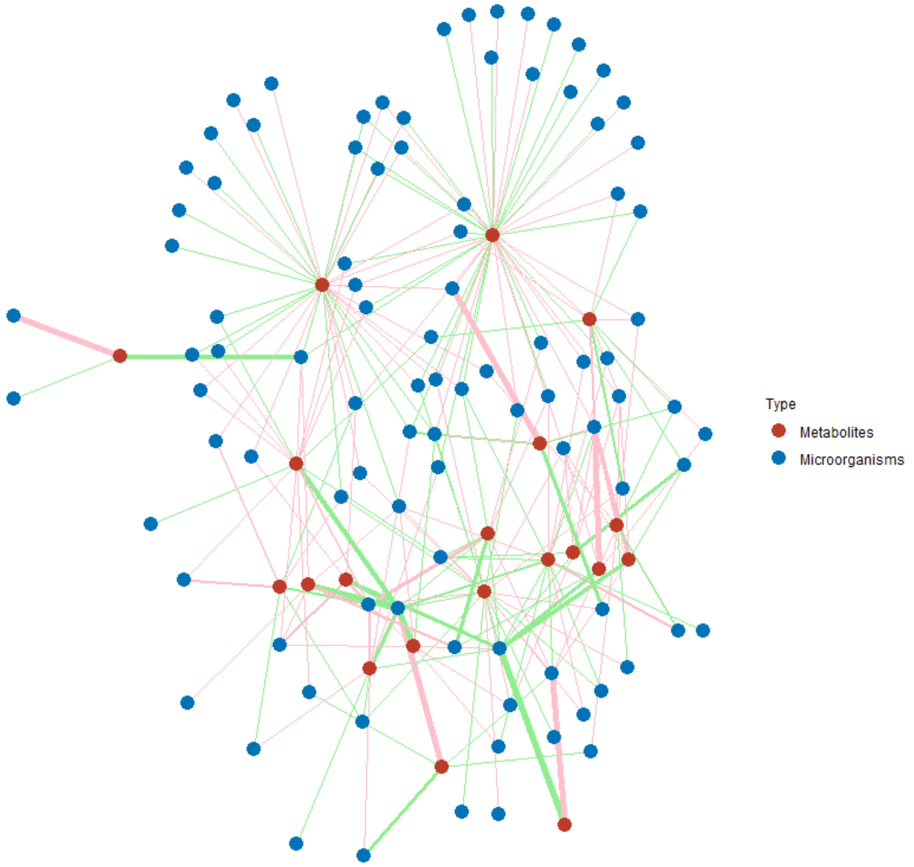

**Figure S17**: Networks of interactions between metabolites (red) and species (blue) provided by the CODA-LASSO for metabolites only found in unaffected samples by sPLS-Reg. Pink edges represent negative associations while green edges represent positive associations.

#### Table

| Method type | Method | Microbiome Normalization | Metabolome Normalization |
|---|---|---|---|
| Global associations | Mantel Test | CLR; ILR; Alpha | Original;Log |
| | MMiRKAT | CLR; ILR; Alpha | Original;Log |
| | Procrustes Analysis | CLR; ILR; Alpha | Original;Log |
| Data summarization | CCA | Original; CLR; ILR; Alpha | Original;Log |
| | PLS-Reg | Original; CLR; ILR; Alpha | Original;Log |
| | PLS-Can | Original; CLR; ILR; Alpha | Original;Log |
| | RDA | Original; CLR; ILR; Alpha | Original;Log |
| | MOFA | Original; CLR; ILR; Alpha | Original;Log |
| Individual Associations | MiRKAT | Original; CLR; ILR; Alpha | Original;Log |

| | | | |
|---|---|---|---|
| | Log-Contrast | Original | Log |
| | Linear Regression/Correlation | CLR | Log |
| | HALLA | CLR | Log |
| Univariate feature selection | CODA-Lasso | Original | Log |
| | clr-(M)LASSO | CLR | Log |
| | (M)LASSO | CLR | Log |
| Multivariate feature selection | sCCA | Original;CLR | Original;Log |
| | sPLS | Original; CLR | Original;Log |

**Table S1: Summary table of the data normalizations used depending on the statistical method**

### Methods

#### Simulation setting

In order to simulate correlated count data from any marginal count distribution we exploited the "Normal to Anything" (NorTA) framework, as already proposed by [1–3]. We provided the general workflow wit the corresponding R code below:

(1) To ensure realistic microbiome and metabolome data we estimated sparse correlation matrices using Spiec-Easi [4] across our three metagenomics-metabolomics datasets. Estimation was performed using the graphical LASSO implemented in the *spiec.easi* function of the *SpiecEasi* R package. As recommended by [4], we estimated the sparse co-occurrence networks of species on the CLR-transformed microbiome to have accurate inferences taking into account the underlying compositionality. Sparse metabolite networks were estimated on the original scale.

```
sparseCov.species = spiec.easi(clr.species, method = "glasso", lambda.min.ratio=1e-2, nlambda=15)$est$cov[[15]]

sparseCov.metabolites = spiec.easi(original.metabolites, method = "glasso", lambda.min.ratio=1e-2, nlambda=15)$est$cov[[15]]
```

(2) Then, we estimated parameters for each microbiota and metabolites, assuming different data generation processes. Parameters were estimated using maximum likelihood. For our

Konzo scenario, we assumed a Negative Binomial distribution for species and a Poisson distribution for metabolites. Similarly, for the Adenomas setting, a zero-inflated negative binomial and a log-normal distribution were assumed for microbiome and metabolome datasets, respectively. Finally, under the Autism scenario, a zero-inflated negative binomial and a Poisson distribution were assumed for microbiome and metabolome datasets respectively. When applicable, Metabolite or Microbiome counts were rounded up to ensure integer. Models specifying only an intercept were fitted to obtain mean, dispersion, and zero-inflation parameters, when required.

***For the Konzo Scenario***:

*glm(Metabolite.Konzo~1, family=poisson())*

*glm.nb(Species.Konzo~1)*

***For the Adenomas Scenario***:

*pscl::zeroinfl(Species.Adenomas~1, dist = "negbin")*

*lm(Metabolite.Adenomas~1)*

***For the Autism Scenario:***

*pscl::zeroinfl(Species.Autism~1, dist="negbin")*

*glm(Metabolites.Autism~1, family =poisson())*

(3) Then, a Gaussian distribution with mean = 0 and a correlation matrix corresponding to the sparse correlation matrices resulting from estimation in (1).

*G1 = MASS::mvrnorm(N, rep(0, n.Metabolites), sparseCov.metabolites )*

*G2 = MASS::mvrnorm(N, rep(0, n.Species), sparseCov.species )*

(4) Data matching the original structures were generated transforming the previous Normal distribution into any arbitrary distribution. This is performed by converting the original Gaussian probability into the quantile of a Negative Binomial, Poisson or Zero-Inflated Negative Binomial distribution, depending on the scenario considered. Parameters estimated in (2) were passed as arguments to ensure synthetic data closely mimic the original data structure. For example, if we want to simulate a correlated zero-inflated Negative Binomial distribution:

***qzinegbin(p = pnorm(G1), size = size.Metabolites, munb = mu.Metabolites ,pstr0 = prop.zeros.Metabolites), ncol=N,nrow=n.Metabolites)***

(5) Under the null hypothesis of no association between metabolites and species, datasets were simulated independently from each other. Thus, any association found in this setup can be considered as a false positive.

(6) Under the alternative, we generated associated microorganisms and metabolites, varying both the number and the strengths of association, as shown in the Figure 2 of the main manuscript.

#### Additional simulation settings

Certain methods require producing additional scenarios in order to match their underlying assumptions.

MMiRKAT presented in the Global association section of the main manuscript requires a smaller number of features than individuals in order to produce results. Under this setting, we randomly subset features of the Adenomas dataset to have half the number of individuals. We generated data under the null and alternative hypothesis.

Similarly, when considering data summarization methods, we generated scenarios with the number of features half the sample size to accommodate assumptions of certain methods, such as CCA which requires N>>P to obtain invertible matrices.

For univariate methods, to reduce the computational burden, we randomly subset features of all datasets to have half their number of individuals. We generated data under the null and alternative hypothesis.

For CODA-LASSO, we produced an additional scenario where we randomly subset 300 species and 600 metabolites. This setting was considered to accommodate long running times of the method.
